# Single-cell atlas of the human immune system reveals sex-specific dynamics of immunosenescence

**DOI:** 10.1101/2024.11.06.622096

**Authors:** Maria Sopena-Rios, Aida Ripoll-Cladellas, Fatemeh Omidi, Sara Ballouz, Jose Alquicira- Hernandez, Roy Oelen, Lude Franke, Monique G.P. van der Wijst, Joseph E. Powell, Marta Melé

**Affiliations:** Life Sciences Department, Barcelona Supercomputing Center (BSC), Barcelona, Spain; Universitat de Barcelona, Barcelona, Spain; Research and Development Department, qGenomics Laboratory, Barcelona, Spain; School of Computer Science and Engineering, University of New South Wales, Sydney, Australia; Center for Data Sciences, Brigham and Women’s Hospital and Harvard Medical School, Boston, MA, USA; Divisions of Rheumatology and Genetics, Department of Medicine, Brigham and Women’s Hospital, Boston, MA, USA; Program in Medical and Population Genetics, Broad Institute of MIT and Harvard, Cambridge, MA, USA; Department of Biomedical Informatics, Harvard Medical School, Boston, MA, USA; Translational Genomics, Garvan Institute of Medical Research, Sydney, NSW, Australia; Department of Genetics, University of Groningen, University Medical Center Groningen, Groningen, the Netherlands; UNSW Cellular Genomics Futures Institute, University of New South Wales, Sydney, Australia

**Author notes:** Contributed equally (Maria Sopena-Rios, Aida Ripoll-Cladellas).

**Keywords:** scRNA-seq, aging, immune system, immunosenescence, sex, females, males, single-cell transcriptomics

## Abstract

Immunosenescence, or immune aging, is characterized by both changes in cell type abundance and a decline in cellular function, leading to increased susceptibility to immune-related diseases. Yet, the extent to which sex influences the dynamic composition of immune cells during immunosenescence remains unknown. Here, we use single-cell RNA sequencing in peripheral blood mononuclear cells from 982 donors to uncover the sex-specific immune aging dynamics. We reveal that aging drives sexually dimorphic compositional and transcriptional shifts, with females showing stronger immune remodeling. Female-specific shifts include the expansion of three cytotoxic CD8+ T effector memory subpopulations and inflammatory monocytes. In addition, female CD4+ central memory T cells exhibit abundance shifts in subpopulations involved in autoimmunity alongside age-associated transcriptional signatures enriched for autoimmune-related pathways. Conversely, males show an age-related expansion of a B cell subpopulation associated with an asymptomatic precursor state to chronic lymphocytic leukemia. These sex-biased, functionally distinct immune subpopulations represent sex-specific hallmarks of immune aging. Our work underscores the complexity and sexual dimorphism of immunosenescence and supports the development of sex-specific strategies to promote healthy aging.

**Highlights:**

‒ A single-cell transcriptomic atlas of >1 million PBMCs from 982 donors uncovers sex-specific immune aging trajectories.
‒ Aging females show greater cellular remodeling and stronger transcriptional shifts than males.
‒ Female-specific expansions with age of cytotoxic CD8+ T effector memory cells, inflammatory monocytes, and CD4+ T helper 22 cells reflect a systemic shift toward a pro-inflammatory, self-reactive immune state.
‒ Male-specific B cell shifts reveal expansion of *CD5+* clonal subpopulations linked to leukemic precursor states.

**Graphical Abstract:** 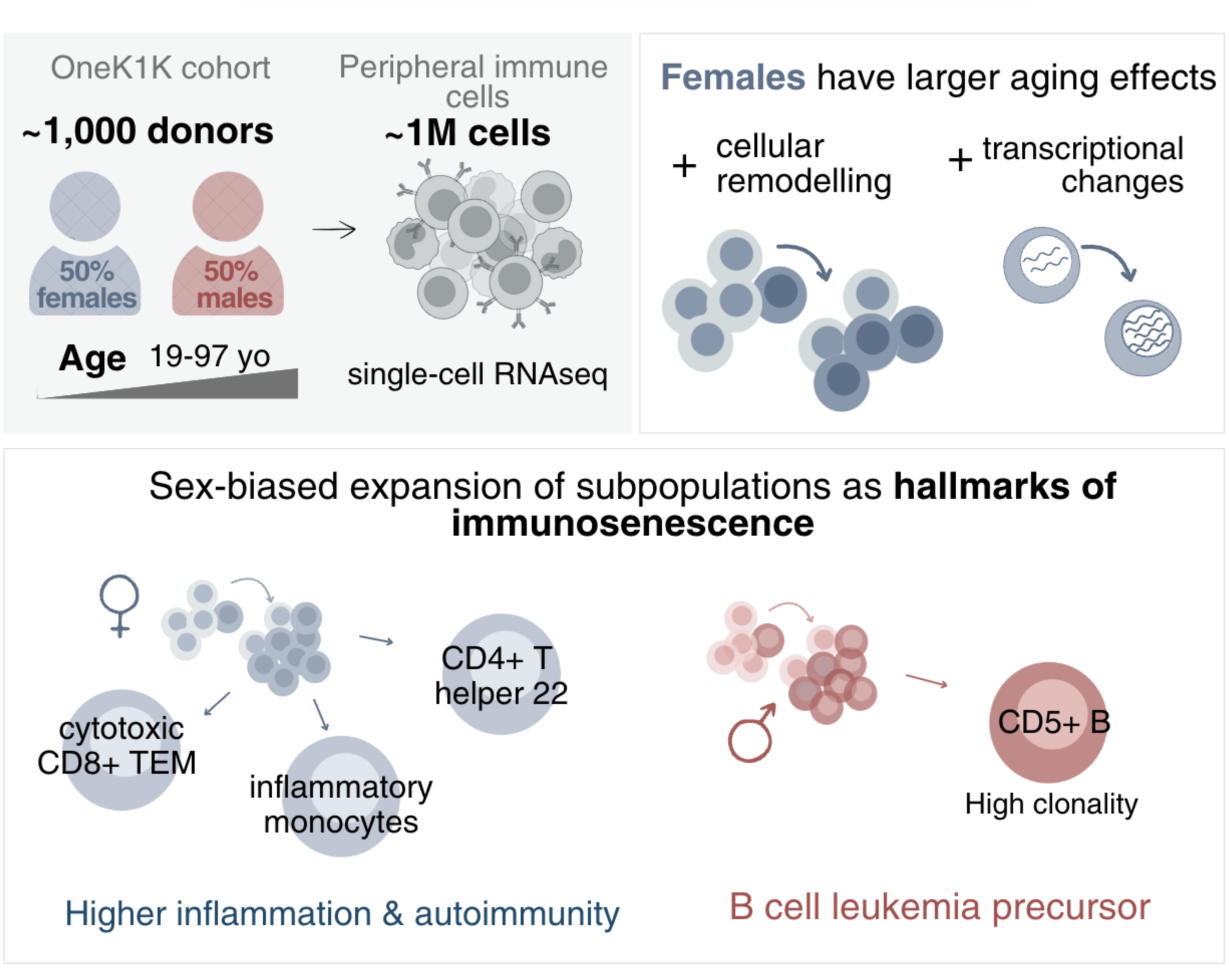

## Introduction

Aging of the immune system, or immunosenescence, refers to the gradual decline of the immune system, which predisposes to multiple diseases, including infection, cancer, autoimmune, and vascular diseases^1,2,3^. This decline hampers the capacity to mount an effective immune response against pathogens and is associated with a persistent low-level inflammatory state, termed inflammaging^4^. Importantly, this process is not uniform across individuals, as males and females exhibit distinct immune responses throughout life. Females typically have stronger immune responses, which confers them greater resistance to infections and improved response to vaccinations^5,6^. However, it also results in 80% higher incidence of autoimmune diseases^7,8^. Aging further introduces sex-specific susceptibilities to other diseases^9,10^, such as cardiovascular^11,12^ or neurological diseases ^13^.

Given the pervasive influence of the immune system throughout the body, immunosenescence also contributes to age-related alterations in other tissues^14,15^. Heterochronic parabiosis experiments, which connect the circulatory systems of young and old mice, show that blood-borne factors can modulate aging across multiple organs, including muscle, liver, brain, and kidney^16–24^. These findings highlight the central role of blood, and by extension the immune system, in systemic aging. As a result, the immune system is emerging as a potential biomarker for aging, serving as a proxy to predict biological age and longevity^25–27^, and may guide the development of healthcare strategies towards healthy aging^28^. Despite this growing recognition, much of immunosenescence research has historically focused on male subjects^29^ and many studies continue to overlook sex differences^30–32^, even as evidence for sexual dimorphism in immune aging continues to accumulate^7,33^.

The age-related decline of the immune system involves both changes in the composition of immune cell populations and molecular alterations^31,32,34–37^. Recent single-cell studies have started elucidating how aging shapes immune cell populations. For instance, aging shifts the balance from naive to more memory T cells^31,38–40^, and certain cytotoxic T cell subtypes become more prevalent^32,34–36^. Moreover, T cells also exhibit the strongest age-associated transcriptional changes among immune cell types^31,41^.

Notably, it has been shown that some age-related molecular changes are sex biased. For instance, males exhibit an age-dependent increase in proinflammatory responses and NK-mediated immunity^34^, accompanied by a more pronounced decline in B-cell specific genes and higher chromatin remodeling^33^. Conversely, females have a more pronounced aging T cell transcriptional signature of increased cytotoxicity and loss of stemness^42^. In addition to these molecular differences, certain immune cell types have been reported to change in abundance with age in a sex-specific manner. For example, elderly males show reduced T cell proliferation and a more modest expansion of CD4+ T memory cells upon antigen re-exposure^43,44^, while young women tend to have higher circulating B cell counts^45^. However, whether specific immune subpopulations undergo sex-specific changes with age is currently unknown. To address this gap, larger and more inclusive studies are needed to explore the sexual dimorphism of the cellular and transcriptional landscape of immune aging, with a focus on specialized subpopulations.

In our study, we provide the largest characterization of the impact of aging on the circulating immune system in over one million peripheral blood mononuclear cells (PBMCs) from 982 young adults to nonagenarians (19-97 years old) across sexes (416 males and 566 females)^46^. We perform both cell type-based and neighborhood-level abundance analyses with age, along with sex-stratified differential gene expression. We reveal that females show greater age-related cellular remodeling and transcriptional changes than males. Moreover, we identify sex-biased immune subpopulations as potential hallmarks of aging, together with a stronger autoimmune transcriptional signature in aging female CD4+ T central memory (TCM) cells. Together, our findings provide a high-resolution map of sex-specific immune aging and lay the groundwork for tailored sex-specific strategies to monitor and improve immune health across the lifespan.

## Results

### Sexually dimorphic trajectories of immune cell composition across the human lifespan

Although aging is known to alter the composition of circulating immune cells^47–50^, whether these compositional changes occur differently in males and females remains unknown. To explore the impact of aging on the circulating immune system at single-cell resolution across males and females, we analyzed single-cell RNA sequencing (scRNA-seq) data of over one million PBMCs from 982 donors from the OneK1K cohort^46^ (**Fig. 1A,B, Fig. S1A,B, Supplementary Table 1**). First, we identified 24 immune cell type populations using Azimuth^51,52^ **(Fig. 1C, D, Fig. S1C).** We then explored age-associated linear shifts in immune cell abundance in each sex separately using a compositional analysis (see **Methods** and **Fig. 1E, Supplementary Table 2**). In both sexes, NK cells increased significantly with age (estimate_females_ = 0.006, FDR_females_ = 0.034; estimate_males_ = 0.008, FDR_males_ = 0.035), consistent with previous findings^32,53,54^. Furthermore, CD8+ T naive cells decreased with age^32,40,41^ and although it was statistically significant only in females (FDR < 0.05), it reached nominal significance in males (p = 0.01) suggesting that CD8+ T decrease with age occurs in both sexes. Conversely, CD8+ T effector memory (TEM) expanded with age exclusively in females (estimate_females_ = 0.009, FDR_females_ = 0.03; estimate_males_ = 0.003, FDR_males_ = 0.64), suggesting that these cells undergo female-specific changes in abundance across the lifespan.

**Figure 1.**
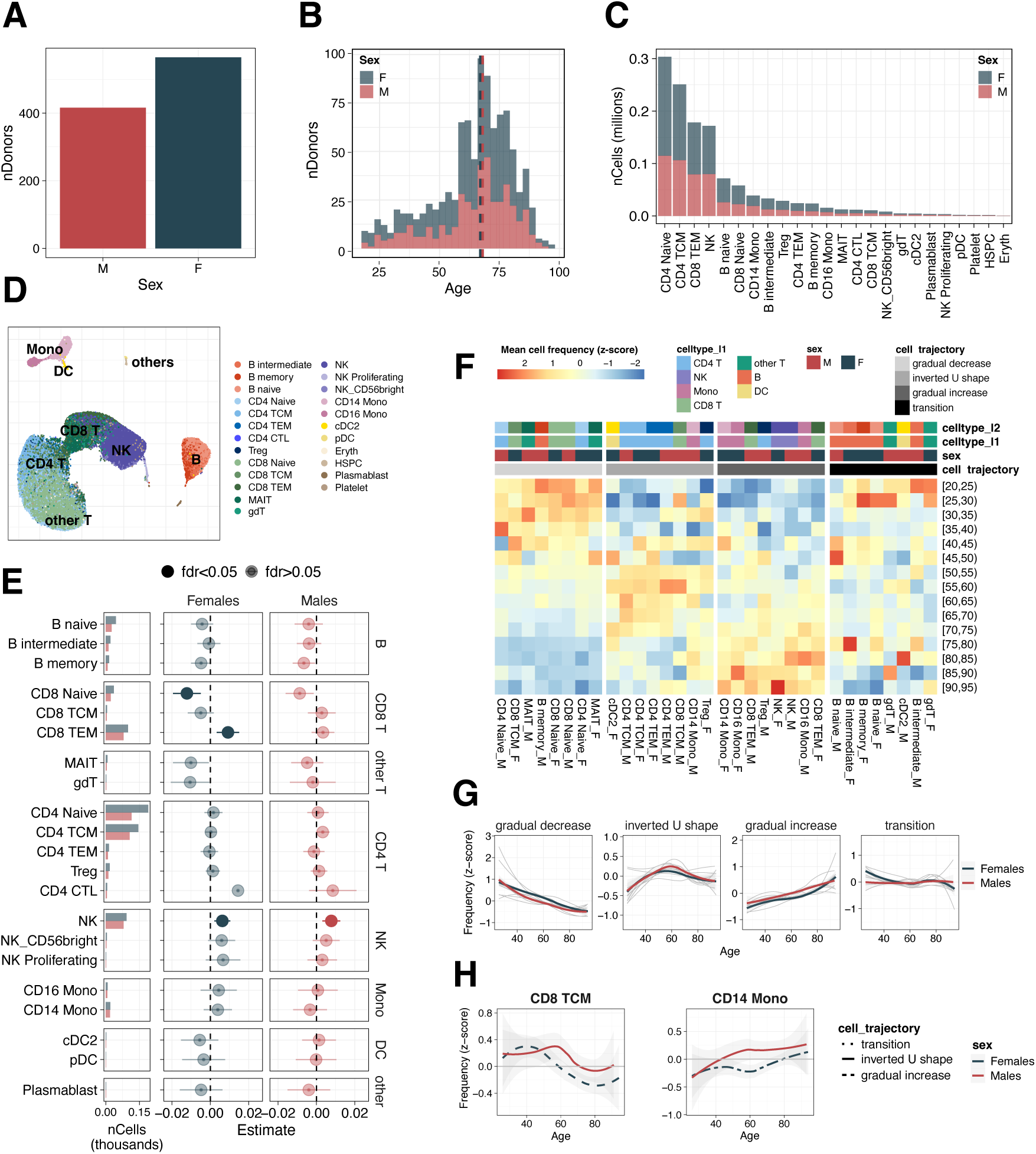
Immune cellular dynamics through lifespan differ between males and females. **A.** Number of donors stratified by sex (416 males (red) and 566 females (blue)). **B.** Age distribution of donors, shown separately for males and females. Vertical dashed lines indicates the median age (males: 68 years old, females: 66 years old) **C.** Total number of cells in each of the 24 immune cell subpopulations annotated using Azimuth^51,52^, stratified by sex **D.** UMAP plot of all immune cells, colored by the 24 annotated immune cell subpopulations. For visualization purposes we downsampled to 500,000 cells (representing ∼50% of the total cells) **E.** Left. Number of cells analyzed per cell type and sex. Right. Estimates from a compositional analysis modeling changes in cell type abundance with age shown separately for females and males. Points represent effect size estimates; bars show 95% confidence intervals. Significant associations (FDR < 0.05) are marked with filled circles. **F.** Heatmap of the normalized mean cell type frequencies (z-scores) across five-year age bins. Rows represent age bins; columns correspond to cell type-sex combinations, ordered by hierarchical clustering. Column annotations indicate cell type (celltype_l2), cell lineage (celltype_l1), sex, and trajectory group. **G.** Distinct aging trajectories of immune cell frequencies across the lifespan. Gray lines represent individual cell types, while colored lines depict the LOESS-smoother mean trajectories for females and males within each trajectory category: gradual decrease, inverted U-shape, gradual increase, and transition. **H.** Representative examples of sexually dimorphic aging trajectories: CD8+ central memory T cells (TCM) and CD14+ monocytes.

Next, we examined whether immune cell populations had non-linear or more subtle aging dynamics and whether these trajectories differed by sex. To address this, we first calculated the mean proportion of each immune cell type in 5-year age bins and clustered them based on their temporal dynamics. We identified four major clusters of immune cells (see **Methods** and **Fig. 1F, G, Fig. S1D, Supplementary Table 2**) with male and female cells of the same cell type generally grouped into the same clusters. The first cluster included cell types gradually declining with age, including naive T cells, a well-documented feature of immune aging^41^. The second cluster followed an inverted U-shape trajectory, expanding in early adulthood before peaking in midlife, and declining thereafter. Most memory T cells, except CD8+ TEM, followed this trajectory. The third cluster comprised populations that progressively increased with age, such as CD8+ TEM, CD16+ monocytes, and NK cells. Notably, these cell types are key mediators of pro-inflammatory response, which may contribute to the persistent or chronic inflammation commonly observed in elderly individuals^50,55^. Finally, the last cluster included cells with fluctuating patterns with B naive and intermediate cells, among others. A few populations had male and female trajectories belonging to different clusters and exhibited distinct sex-specific temporal dynamics (**Fig. 1H, Fig. S1D, E**). For instance, CD8+ TCM cells declined more steeply and earlier in females, while CD14+ monocytes increased earlier in males. In addition, although both male and female CD8+ TEM cells belonged to the gradual increase cluster, young CD8+ TEM were less abundant in females (estimate_<50_= 0.29, p-value_<50_=0.048, estimate_≥50_=-0.02, p-value_≥50_=0.77) and experienced a steeper increase in abundance with age compared to males. These differences suggest that some immune trajectories may be modulated by sex-specific physiological changes over the lifespan.

Overall, our findings indicate that sex-specific differences can shape immune cell type trajectories, underscoring the complex interplay between aging, immune composition, and sex-specific regulation.

### Neighborhood-level analysis reveals larger immune cell remodeling with age in females

Our previous analysis identified cell types that exhibit age-related changes in abundance that differ between males and females. However, this analysis is limited to annotated cell types and may therefore disregard more subtle, sex-specific abundance changes occurring only in specific cell subpopulations^32,36,56^. To address this, we used Milo, a framework for differential abundance testing that does not rely on discrete cell type labels^57,58^. First, we constructed a k-nearest neighbor (KNN) graph where each node represents a cell, and the edges connect each cell to its nearest neighbors based on gene expression similarity. From this graph, we defined cellular states as neighborhoods with overlapping cells, which were then used for differential abundance testing with age separately in each sex. Lastly, we mapped each neighborhood to its corresponding pre-annotated cell type to enhance interpretation and compare our cell type-agnostic approach to the previous cell type-based composition analysis (see **Methods** and **Fig. 2A, Fig. S2A, B, Supplementary Table 3**). Both approaches are highly concordant (Spearman R_females_ = 0.82, p-value_females_ = 0.001; Spearman R_males_ = 0.71, p-value_females_ = 0.002) **(Fig. S2C)**, but the neighborhood-based approach has greater statistical power to detect subcellular changes. Using this approach, we identified neighborhood-level compositional changes in almost all cell types (in 13 cell types in females and in 12 cell types in males).

**Figure 2.**
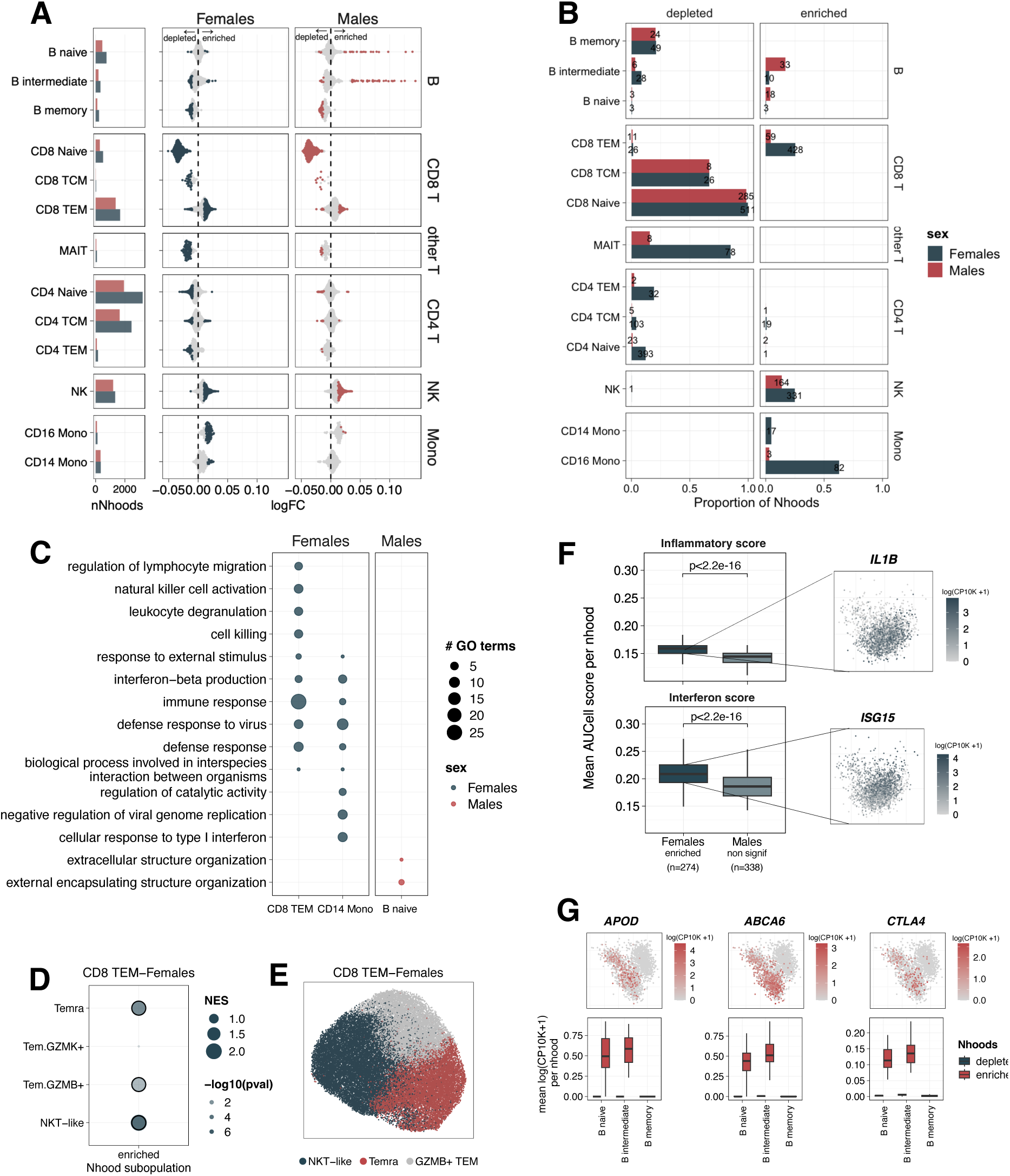
Cell-type–agnostic analysis reveals sex-specific expansion of functionally distinct immune subpopulations with age. **A.** Left. Number of neighborhoods tested per cell type, stratified by sex. Right. Beeswarm plot showing the distribution of age-associated log fold changes (logFC) in neighborhood abundance for females (blue) and males (red). Each dot corresponds to a neighborhood of cells assigned to a dominant cell type (>80% of cells within the neighborhood (see **Methods**)). Neighborhoods significantly associated with age (Spatial-FDR < 0.05) are colored. **B.** Proportion and number of age-associated neighborhoods that are significantly depleted (left) or enriched (right), shown by cell type and sex. **C.** Non-redundant set of Gene Ontology (GO) terms (FDR < 0.05) enriched among marker genes of age-enriched neighborhoods in females and males. **D.** Gene Set Enrichment Analysis (GSEA) of marker genes from female age-enriched CD8+ T effector memory (TEM) neighborhoods, compared to transcriptional signatures of CD8+ TEM subpopulations (Temra, *GZMK+* TEM, *GZMB+* TEM, and NKT-like) from^32^. Significant enrichments (FDR < 0.05) are highlighted. Dot size reflects the normalized enrichment score (NES). **E.** UMAP plot of cells from female age-enriched CD8+ TEM neighborhoods, colored by clusters corresponding to three cytotoxic subpopulations: NKT-like, Temra and *GZMB+* TEM. **F.** Left. Distribution of inflammatory (top) and interferon scores (bottom) for female age-enriched CD14+ monocyte neighborhoods compared to male non-significant CD14+ monocyte neighborhoods. Right. UMAP of female-age-enriched CD14+ monocyte cells colored by the expression of a top gene marker related to inflammation (*IL1B,* top) and an interferon-stimulated gene (*ISG15,* bottom). **G.** Top. UMAP plots of male-age-enriched B naive and intermediate cells colored by expression of top marker genes (*APOD, ABCA6,* and *CTLA4*). Bottom. Average log-normalized, scaled expression of the same genes across age-enriched or age-depleted B cell neighborhoods in males.

In females, 17% of neighborhoods change abundance with age compared to 7% in males, suggesting that immune cells undergo comparatively larger abundance shifts with age in females (**Fig. 2B**). These sex-differences relate to both the proportion of neighborhoods that increase (11% in females vs 5% in males) and decrease in abundance with age (8% in females and 3% in males). The largest sex-difference occurs in MAIT cells in which females have an age-related decline of nearly the entire population (85% of neighborhoods) whereas in males only 16% of neighborhoods decline. Females also have a larger proportion of neighborhoods increasing with age in CD16+ Monocytes (63% in females vs. 3% in males), CD14+ Monocytes (5% in females vs. 0% in males) and in CD8+ TEM (25% in females vs. 4% in males) consistent with our sex-stratified compositional analysis **(Fig. 1A)**. In contrast, B naive and B intermediate cells have more neighborhoods increasing abundance with age in males (B naive: 0.039% in males vs. 0.004% in females; B intermediate: 0.17% in males vs. 0.03% in females). Interestingly, these male-specific shifts have the largest fold changes across all the cell types (mean-log fold change (logFC) = 0.06, and B intermediate, mean-logFC = 0.07) **(Fig. 2A)**.

In summary, these findings demonstrate that most immune cell types undergo subpopulation abundance shifts with age and suggest that these are more pronounced in females.

### Cytotoxic CD8+ TEM and *CD5+* B cell subpopulations represent sex-specific hallmarks of aging

Changes in neighborhood abundance with age may reflect shifts of functionally distinct cellular subpopulations^57–60^ that could have downstream consequences for the immune system’s function. To further explore whether age-associated neighborhoods represent immune subpopulations with distinct biological functions, we identified marker genes for these neighborhoods within each cell type and performed pathway enrichment analysis (see **Methods**). We identified three sex-specific, age-enriched neighborhood groups —in CD8+ TEM and CD14+ Mono cells in females and B naive cells in males— with pathways enriched in particular biological functions **(Fig. 2C, Supplementary Table 3)**.

In females, CD8+ TEM neighborhoods accumulating with age have gene expression signatures related to cell killing and NK activation^56^ **(Fig. 2C, Fig. S2D)**, suggesting a cytotoxic phenotype^61^. A key gene of this cytotoxic-related signature is Granzyme B (*GZMB*) **(Fig. S2D)**, a gene previously reported to be a marker gene of three types of CD8+ TEM subpopulations: NKT-like, *GZMB*+ TEM, and terminally differentiated CD45RA+ memory (Temra) cells^32^. To determine which of these subtypes our age-enriched neighborhoods correspond to, we performed gene signature enrichment and found significant overlap with all three (see **Methods** and **Fig. 2D, Supplementary Table 3**) (NES_NKT-like_ = 2.07, FDR_NKT-like_ = 5.4·10^-08^, NES_GZMB+_ = 1.58, FDR_GZMB+_ = 3.87·10^-03^, NES_Temra_ = 1.90, FDR_Temra_ = 9.65·10^-06^).

Unsupervised clustering of cells within these *GZMB-*expressing effector memory neighborhoods confirmed this result: each of the three identified clusters expressed markers corresponding to one of the *GZMB*-expressing CD8+ TEM subtypes^32^ (see **Methods** and **Fig. 2E, Fig. S2E)**. Notably, when performing sex-stratified age-compositional analysis separating between *GZMB*+ and *GZMB*-CD8+ TEM cells (see **Methods** and **Supplementary Table 3**), only female *GZMB*+ cells increased with age (estimate_females_ = 0.02, FDR_females_ = 1.1×10⁻³; estimate_males_ = 0.007, FDR_males_ = 0.3) **(Fig. S2F)**. This suggests that the increase in *GZMB*-expressing effector memory subpopulations drives the broader CD8+ TEM expansion in aging females observed in previous analysis **(Fig. 1A)**. We replicated this finding in a smaller dataset^62^ (estimate_females_ = 0.05, p-value_females_ = 0.01) (see **Methods** and **Supplementary Table 3**). Together, these results showed that all three cytotoxic CD8+ TEM subpopulations (NKT-like, GZMB+, and Temra) accumulate with age specifically in females and may collectively underlie the observed female-specific expansion of CD8+ TEM cells.

The CD14+ Monocyte subpopulation that expands with age in females was enriched for genes involved in viral defense and type I interferon (IFN) signaling **(Fig. 2C)**. This suggests a heightened inflammatory and IFN-driven response in the aging female myeloid compartment, potentially contributing to inflammaging^63,64^. Consistent with this observation, these cells also showed increased expression of IFN-stimulated genes (estimate = 0.018, FDR = 7.3·10^-03^) and inflammatory genes (estimate = 0.019, FDR = 9.3·10^-12^) (see **Methods** and **Fig. 2F, Supplementary Table 3)**.

Finally, in males, B naive cell neighborhoods enriched with age expressed marker genes related to lipid-transport, including *ABCA6* and *APOD* **(Fig. 2G),** which matched the transcriptional signature of a previously characterized *CD5+* B cell population^32,36,56^ **(Fig. S2G,H, Supplementary Table 3).** Indeed, we replicated this male-specific accumulation of *CD5+* B cells with age in males in a smaller dataset^32^ (estimate_males_ = 0.26, FDR_males_ = 0.0031). Marker genes of age-accumulating neighborhoods of B intermediate cells in males were also enriched for *CD5+* B cell signatures **(Fig. 2G, Fig. S2H,I, Supplementary Table 3).** Notably, over 40% of the cells in male age-enriched *CD5+* B cell neighborhoods originated from only two donors **(Fig. S2J-L)**, suggesting that *CD5+* expansions may be largely individual-specific (see **Methods** and **Supplementary Table 3**). While *CD5*+ B cell subpopulation is known to have the most prominent clonal expansion with age^32^, it remains unclear whether this age-related increase in clonality differ between males and females^32^. Upon sex-stratified re-analysis of a BCR-sequencing dataset^32^, we show that *CD5+* B cell clonal expansions with age occur predominantly in males (estimate_males_ = 0.17, FDR_males_ = 4.27·10^-04^; estimate_females_ = 0.009, FDR_females_ = 0.7) **(Fig. S2M),** uncovering a previously unrecognized male bias. This is particularly relevant given that expanded *CD5+* B cell clones are associated with monoclonal B-cell lymphocytosis, an asymptomatic precursor of chronic lymphocytic leukemia, both more prevalent in males^65–67^. Thus, the male-biased expansion of *CD5+* B cells may reflect early immune aging processes that increase susceptibility to B cell malignancies in aging males.

Overall, by employing a cell type-agnostic approach, we identify, for the first time, sex-specific subpopulations that accumulate with age —insights that may be overlooked by traditional cell type annotations— and that may represent hallmarks of the aging immune system^13,14,71^.

### Stronger transcriptional responses in females underlie sex-biased immune aging

Sex differences in age-related immune decline might be shaped not only by shifts in cell type composition but also by transcriptional changes across immune cell types^34^. To address this, we performed differential gene expression analysis with age stratifying by sex, controlling for technical (sequencing pool and number of cells per donor) covariates (see **Methods**). We identified age-differentially expressed genes (age-DEGs) in almost all tested cell types across both sexes (19 out of 24 immune cell types, 79%) (FDR< 0.05) **(Fig. 3A, Supplementary Table 4).** Males and females shared around 25% of age-DEGs across cell types, indicating a shared aging signature in immune cells between sexes (**Fig. S3A**). T cells had the largest number of age-DEGs, followed by B and NK cells, consistent with the well-established remodeling of the T cell compartment as a hallmark of immune aging^31,41,68,69^, a pattern that held regardless of donor sample size (see **Methods** and **Fig. S3B, Supplementary Table 4**). Notably, females had a substantially higher number of age-DEGs across most cell types (2,306 age-DEGs in females and 1,122 in males adding up all cell types). The most significant sex-biased cell types were CD8+ TEM and NK cells, where females had 10-fold and 11-fold more age-DEGs than males, respectively (Binomial test, FDR_CD8TEM_ = 1.21·10^-85^, FDR_NK_ = 1.65·10^-53^) (**Fig. S3C**). This female-biased response persisted both after downsampling to equal donor numbers between sexes and when testing the age-by-sex interaction, despite the lower statistical power (see **Methods** and **Fig. S3D, E, Supplementary Table 4**). Overall, these results suggest that females have a stronger transcriptional response to aging.

**Figure 3.**
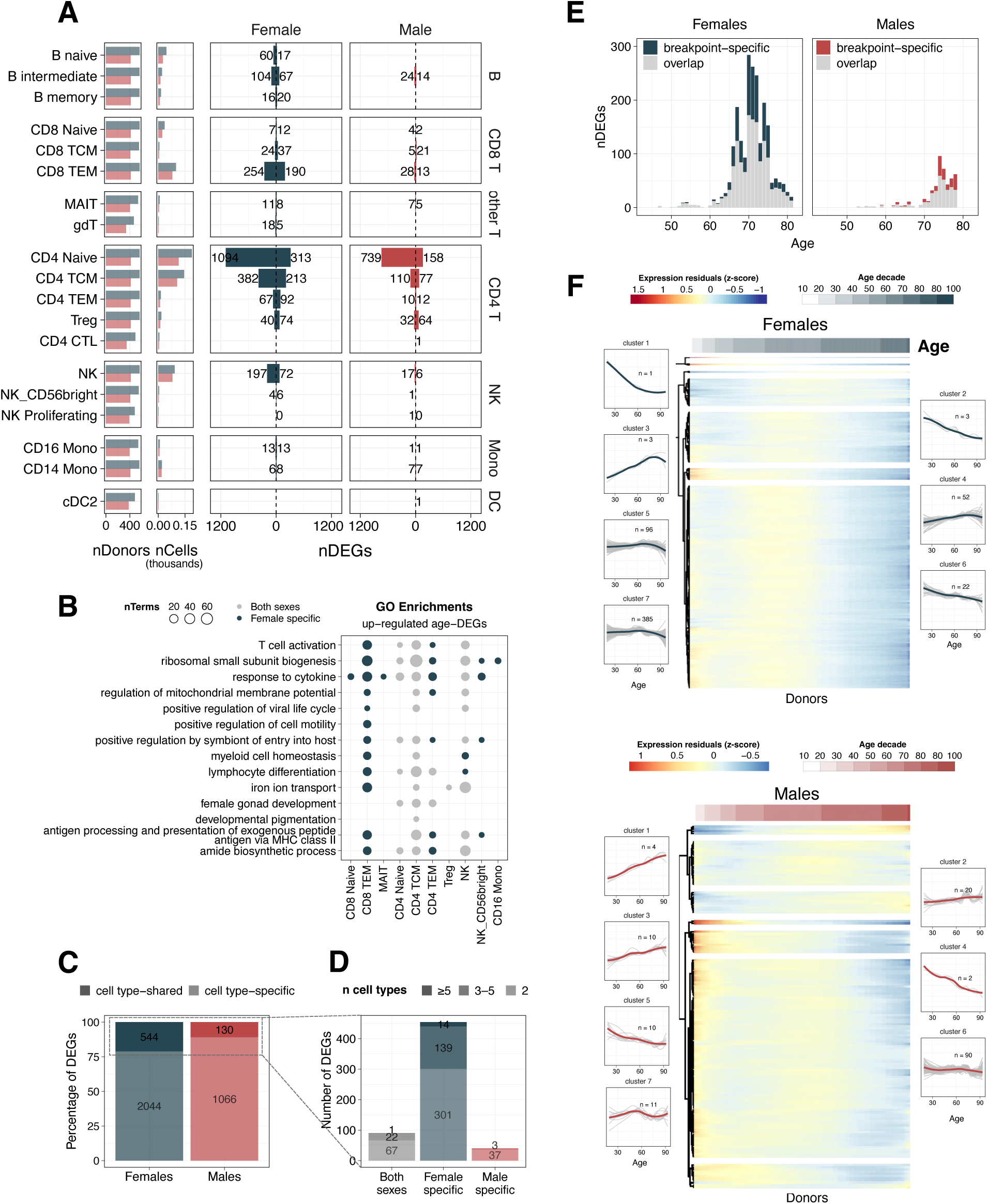
Females exhibit more extensive age-associated gene expression changes across immune cell types. **A.** Left. Number of donors analyzed per cell type and sex. Center. Number of cells analyzed per cell type and sex. Right. Number of age-differentially expressed genes (age-DEGs; FDR < 0.05) per immune cell type in females (blue) and males (red). **B.** Non-redundant set of Gene Ontology (GO) terms enriched among up-regulated age-DEGs, highlighting biological pathways associated with sex-shared versus sex-specific aging signatures. **C.** Percentage and total number of age-DEGs that are cell type-specific versus shared across multiple cell types, shown separately for females and males. **D**. Number of cell type-shared age-DEGs, grouped by the number of cell types in which the gene is differentially expressed: low (2 cell types), intermediate (3–5 cell types), and high (≥5 cell types), stratified by female-specific, male-specific, and sex-shared signatures. **E.** Number of age-DEGs detected in CD4+ T naive cells using a 30-year sliding window approach (“breakpoint analysis”; see **Methods**), colored by genes uniquely identified through the breakpoint analysis and those overlapping with the standard linear model. **F.** Heatmap of residual z-scored gene expression trajectories in CD4+ T naive cells, based on LOESS-smoothed expression profiles of age-DEGs identified through breakpoint analysis and grouped by hierarchical clustering (see **Methods**). Top: females; bottom: males. Adjacent line plots display the average expression trajectory of genes within each of the seven clusters, with bold lines representing the smoothed mean.

Functional enrichment analysis on age-DEGs identified significant enrichments in 13 immune cell types, with a total of 20 enriched pathways **(Fig. 3B, Fig. S3F** and **Supplementary Table 4**). Among them, approximately half were shared between sexes, while the rest were female specific. These pathways spanned immune functions such as T cell activation and cytokine responses, as well as metabolic processes and mitochondrial activity. Despite a strong female bias in the number of age-DEGs in NK cells **(Fig. 3A, Fig. S3C)**, the affected biological pathways were largely shared between sexes, indicating similar age-related molecular changes **(Fig. 3B, Fig. S3F)**. In contrast, while CD8+ TEM cells had a similarly strong female bias in the number of age-DEGs, the functional enrichments were significant only in females, indicating female-specific age-related remodeling. This female bias is likely driven, in part, by the age-related expansion of the *GZMB*+ CD8+ TEM cell subpopulation observed especially in females. Furthermore, although most age-DEGs were cell type-specific (79% in females and 89% in males)^41^ **(Fig. 3C,D),** many of the underlying biological pathways were shared across cell types, suggesting a broad, orchestrated response to aging **(Fig. 3B)**.

Gene expression changes in the immune system across the lifespan can follow non-linear trends, with certain temporal changes occurring more rapidly at specific ages^33,70–72^. To investigate this, we stratified individuals into three overlapping age groups —early (<50 years old), mid (40–60 years old), and late (≥50 years old)— and performed differential expression analysis with age within each age group and sex (see **Methods** and **Fig. S4A, Supplementary Table 5**). Most transcriptional changes occurred in the late-age group in both males and females. However, downsampling analysis indicated that sample sizes in the early and mid-groups were likely insufficient to capture age-related shifts at earlier life stages **(Fig. S4B)**. Next, we wanted to identify whether these observed late-related changes in gene expression occurred at particular time points in the adult life^33,70,71^. To this end, we reimplemented the sliding-window approach described in^71^, referred to here as breakpoint analysis, comparing consecutive 15-year age groups (e.g., 35–50 vs. 50–65 years old), shifting the 30-year window by one year across the age range (see **Methods** and **Supplementary Table 5**). Consistent with our previous findings **(Fig. S4A)**, we observed a late-life peak in gene expression changes, most prominent in CD4+ T naive cells, the most abundant cell type **(Fig. 3E)**. This peak occurs around age 70 in females and five-years later and subtler in males. Despite reduced statistical power, similar female-specific peaks were also detected in less abundant cell types (such as CD4+ TCM, NK, and B naive cells) **(Fig. S4C, D)**, supporting a coordinated but sex-dependent pattern of immune aging. Notably, approximately half of the genes identified in the breakpoint analysis overlapped with those from standard linear differential gene expression analysis (mean_females_ = 57.2%, mean_males_ = 48%) **(Fig. S4E, F)**, suggesting that linear models, while not designed to detect inflection points, can still capture abrupt transcriptional change. To further characterize the wave-like dynamics in CD4+ T naive cells, we grouped genes with similar trajectories using unsupervised hierarchical clustering and identified many clusters with non-linear trajectories (see **Methods** and **Fig. 3F** and **Supplementary Table 5**). Several of these clusters were enriched for functional pathways (FDR < 0.05) —including signaling and RNA binding in females, and cytokine and growth factor binding in males— further supporting distinct, yet coordinated, shifts in biological processes during aging **(Fig. S4G** and **Supplementary Table 5)**.

Taken together, our results highlight a broader and stronger female transcriptional response, especially in immune effector cells such as CD8+ TEM and NK cells, which may be accompanied by abrupt gene expression changes at specific stages of the adult lifespan.

### Female CD4+ TCM cells exhibit increased autoimmune-related expression and abundance shifts with age

Sex differences in immune aging may help explain why many age-related diseases exhibit sex-biased susceptibility^30,73^. To investigate this, we performed functional enrichment of male and female age-DEGs using DisGeNET, a comprehensive gene–disease association database^74,75^ (see **Methods** and **Fig. 4A, Supplementary Table 6**). Most enriched disease terms (25 of 32, 78%) were primarily associated to CD4+ TCM cells Notably, 83% of CD4+ TCM-enriched terms were autoimmune-related, and four of them were uniquely enriched in female-specific age-DEGs, consistent with the important role of CD4+ TCM in autoimmune disease pathogenesis^76–78^. To test whether these age-transcriptional shifts in CD4+ TCM cells could reflect changes in autoimmune potential, we computed an autoimmunity score per cell based on the enrichment of an independent, manually curated autoimmune disease gene set^79,80^ (see **Methods** and **Supplementary Table 6)**. Autoimmunity scores increased significantly with age in both sexes (estimate_males_ = 2.33·10^-5^, FDR_males_= 0.022; estimate_females_ = 2.85·10^-3^, FDR_females_=0.021) **(Fig. 4B)**. However, females exhibited significantly higher scores after the age of 50 compared to males (Wilcoxon rank-sum test (under age 50) p = 0.64, (over age 50), p = 0.0026), suggesting that upon aging, females have a stronger shift toward a more self-reactive state.

**Figure 4.**
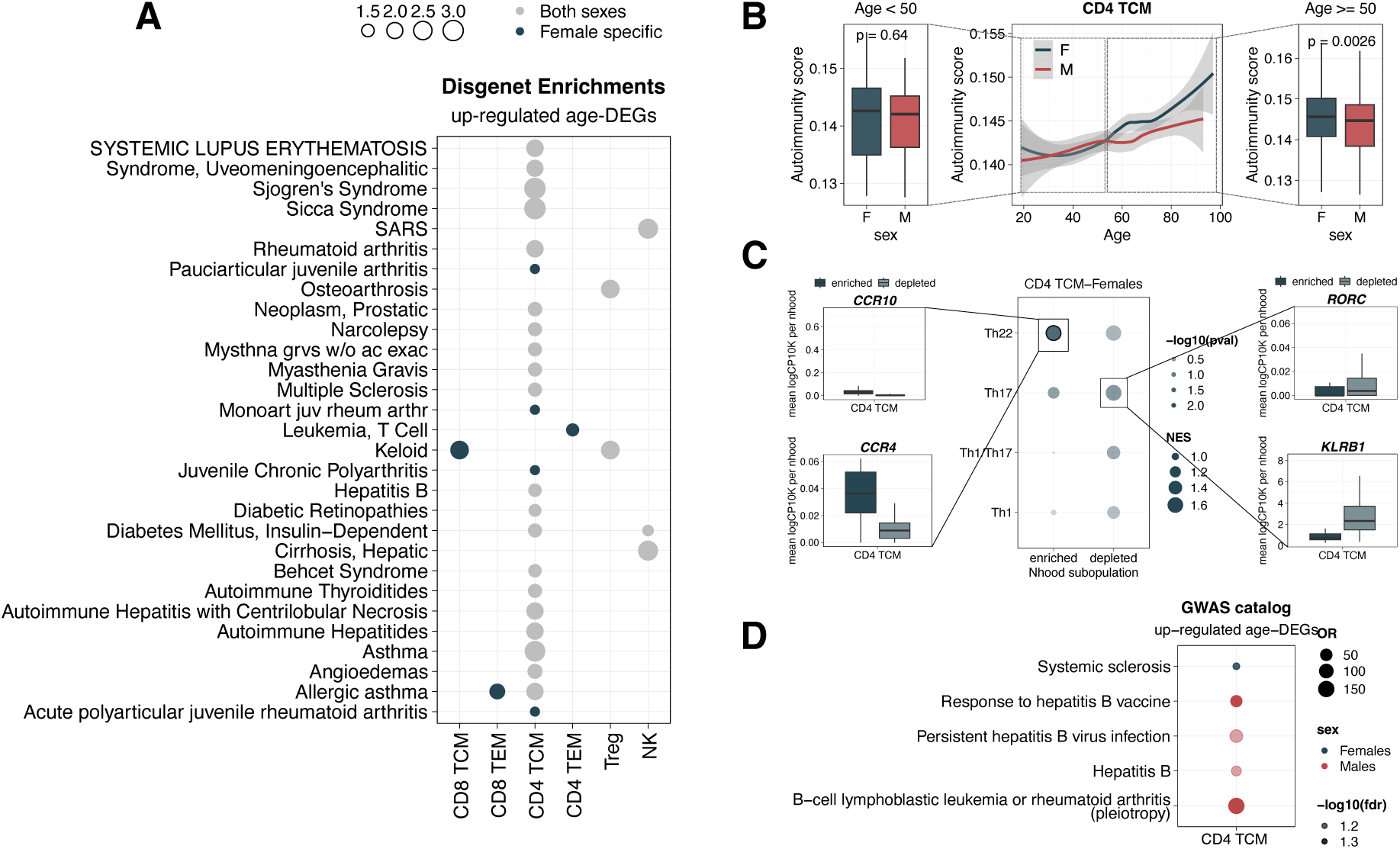
CD4+ TCM cells in females exhibit age-driven autoimmune-related expression and abundance changes. **A.** Disease enrichment analysis using DisGeNET^74^, showing associations between age-DEGs (sex-shared and sex-specific) and gene sets linked to known diseases. **B** Left. Distribution of the donor-level mean autoimmunity score in CD4+ T central memory (TCM) cells for females (blue) and males (red) under age 50. Center. Age-related trajectories of donor-level mean autoimmunity scores in CD4+ TCM cells, shown separately by sex. Right. Distribution of the donor-level mean autoimmunity scores in individuals over age 50. **C.** Left. Average log-normalized, scaled expression of CD4+ T helper 22 (Th22) marker genes (*CCR4, CCR10*) across neighborhoods. Center. Gene Set Enrichment Analysis (GSEA) of marker genes from age-enriched CD4+ TCM neighborhoods in females, compared to transcriptional signatures of CD4+ TCM subpopulations (Th22, Th17, Th1/Th17, and Th1) from^32^. Significant enrichments (FDR < 0.05) are highlighted. Dot size reflects the normalized enrichment score (NES). Right. Average log-normalized, scaled expression of CD4+ T helper 17 (Th17) marker genes (*CCR4, CCR10*) across neighborhoods. **D.** Overlap between sex-stratified age-DEGs and Genome-Wide Association Studies (GWAS) hits (FDR < 0.1). Dot size reflects the odds ratio (OR) from a Fisher’s exact test assessing the significance of the overlap.

We next wondered whether abundance changes within CD4+ TCM cell subpopulations could contribute to the observed increase of autoimmune-related signatures with age. Most aging-related subpopulation abundance changes in CD4+ TCM cells occurred in females (122 neighborhoods changed abundance in females vs. 6 in males) **(Fig. 2A, B).** To characterize these age-associated CD4+ TCM neighborhoods, we performed gene signature enrichment using marker genes from previously described CD4+ TCM subpopulations^32^ (see **Methods** and **Fig. 4C, Supplementary Table 6**). Both age-enriched and age-depleted female neighborhoods showed transcriptional profiles matching well-defined T helper (Th) subpopulations^32,81,82^. Specifically, enriched neighborhoods were mainly associated with CD4+ Th22 cells (NES = 1.42, FDR = 0.03), whereas depleted neighborhoods belonged to CD4+ Th17 cells (NES = 1.79, FDR = 0.19) which are known to decline with age^83^. Both Th17 and Th22 subpopulations are well-recognized contributors to autoimmune pathogenesis^84–88^. Th17 cells drive inflammation through IL-17 secretion, promoting neutrophil recruitment and the production of pro-inflammatory cytokines and chemokines^89^. Th22 cells, a more recently defined subset^81,90,91^, secrete IL-22 and contribute to tissue remodeling and inflammation, particularly in epithelial barriers^81^. Thus, females undergo stronger abundance changes of Th22 and Th17 cell subpopulations that may heighten autoimmune susceptibility and progression with age.

Finally, given the strong heritability of autoimmune diseases^92^, we examined whether age-DEGs were enriched for disease-associated genetic variants in the Genome-Wide Association Studies (GWAS) catalog^93^ (**see Methods** and **Fig. 4D, Supplementary Table 6**). Only CD4+ TCM cells showed significant enrichments (FDR < 0.1). In females, age-up-regulated DEGs were enriched for one autoimmune disease, systemic sclerosis (OR = 7.81, FDR = 0.059), consistent with previous findings showing that menopause-associated estrogen loss can worsen systemic sclerosis symptoms^94,95^. In contrast, in males, enriched traits included mostly hepatitis B virus (HBV)-related terms (mean OR = 49.88, FDR = 0.075), and pleiotropic genes linked to both B-cell lymphoblastic leukemia and rheumatoid arthritis (OR = 150, FDR = 0.059), consistent with known male biases in infection outcomes and non-reproductive cancer risk^7,96,97^, as well as elderly-onset rheumatoid arthritis^98^.

Together, these results reveal novel associations between sex-specific changes with age and disease susceptibility and suggest that transcriptional and compositional remodeling within CD4+ TCM cells contributes to the increased autoimmune potential observed in aging females.

## Discussion

Males and females differ in their innate and adaptive immune responses, contributing to sex-biased disease susceptibility. However, how sex shapes immune aging —a key driver of age-related disease— remains poorly understood^30^. Here, we provide the most comprehensive single-cell characterization to date of sex-specific immune aging dynamics in humans. We show that immune aging is sexually dimorphic, with females exhibiting more pronounced cellular remodeling and stronger transcriptional responses than males. These differences likely reflect the cumulative influence of genetic, hormonal, and environmental factors —including sex chromosome-linked genes, reproductive status, and exposures such as nutrition and the microbiome— which interact over time to shape immune trajectories^7^. Crucially, we uncover sex-specific immune subpopulations that expand with age and display distinct functional phenotypes. These populations may serve as early hallmarks of immunosenescence and could represent mechanistic precursors to the divergent disease vulnerabilities observed in aging males and females.

Females typically mount stronger immune responses, which improve resistance to infections and vaccine efficacy^5,6^, but also result in an 80% higher incidence of autoimmune diseases^7,8^. Aging itself is an additional risk factor for autoimmune diseases^99,100^, yet, the relationship between aging and autoimmunity vulnerability in females remains complex^101,102^. In some autoimmune diseases, menopause and its associated drop in estrogen worsen disease severity —as observed in multiple sclerosis^103,104^ and rheumatoid arthritis^105,106^ — while in others, such as systemic lupus erythematosus, disease activity may improve with age^107^. Here, we show that specific cell subpopulations with pivotal roles in autoimmune diseases increase in abundance with age especially in females, providing new cellular and molecular insights into these patterns. For instance, aging in females is accompanied by the expansion of cytotoxic *GZMB-*expressing CD8+ T cell subpopulations **(Fig. 2D, E),** which have been recently identified as primary drivers of the IFN signature in five autoimmune conditions, including rheumatoid arthritis, psoriatic arthritis, ulcerative colitis, Crohn’s disease, psoriasis^108^. Similarly, the accumulation of inflammatory monocytes in aging females **(Fig. 2F)** is a recognized hallmark of several autoimmune diseases, such as rheumatoid arthritis^109,110^ and inflammatory bowel disease^111,112^. Together, these expansions may reflect a systemic shift toward a more cytotoxic, pro-inflammatory, autoimmune-like state in aging females. Finally, we observe a stronger autoimmune signature in aging female CD4+ TCM cells **(Fig. 4)**, including up-regulation of autoimmune-related genes, increased autoimmunity scores after midlife, and a selective expansion of Th22 cells. Th22 cells, which secrete IL-22 and promote inflammation^81^, have been implicated in a growing number of autoimmune conditions^84–88^, suggesting that their age-related accumulation may exacerbate late-life autoimmunity in females. These findings may help explain why older females are more susceptible to chronic inflammatory and autoimmune diseases, despite stronger immune surveillance. Monitoring the expansion of these sex-biased subpopulations could enable earlier detection of age-related autoimmune alterations, as well as guide targeted, sex-specific interventions to prevent or mitigate late-life immunopathologies.

In contrast to females, males are more vulnerable to certain malignancies, showing higher incidence and mortality rates for most cancer types^96,113,114^. This excess persists —and in some cases widens— into late life, suggesting that sex differences in cancer risk are not solely due to the protective effects of female hormones before menopause^113,114^, but also reflect broader age-related biological and behavioral factors^114,115^. Our study shows that, overall, immune aging in males is less transcriptionally pronounced, with fewer sex-specific signatures and more modest shifts in immune cell abundance. However, we observe individual-specific expansions in *CD5+* B cells, a subpopulation marked by increased clonality and lipid-related gene expression **(Fig. 2G, Fig. S2G-M)**. The age-related expansion of this subpopulation may represent early stages of monoclonal B-cell lymphocytosis, a precursor to chronic lymphocytic leukemia, which is more prevalent in older males^67^. Hence, our findings provide molecular evidence that male-biased expansion of clonal *CD5+* B cells may contribute to the elevated risk of B cell malignancies in aging males^116^. Consistently, we find enrichment of age-associated genes in CD4+ TCM cells at B-cell lymphoblastic leukemia loci^117^ **(Fig. 4D)**. Beyond cancer, males and females also differ in infection prevalence, severity, and pathogenesis, with males generally exhibiting greater susceptibility to infection^118^. In this line, we show that older males increase expression of genes associated with hepatitis B (HBV) susceptibility **(Fig. 4D)**, mirroring the higher HBV viral load in aging males^7,119–122^. Chronic HBV infection is also a known risk factor for hepatocellular carcinoma^123^, a malignancy that tends to be more prevalent in older men^124^. Altogether, our findings illustrate how male-specific immune aging patterns may contribute to increased vulnerability to hematologic malignancies and chronic infections. These shifts could serve as risk biomarkers and inform sex-specific preventive strategies.

Finally, the sex-specific immune subpopulations we identify may serve as hallmarks of immunosenescence that reflect —and potentially drive— immune remodeling across peripheral tissues. Immunosenescence arises through both intrinsic aging of immune cells and extrinsic effects of tissue senescence, which compromise barrier integrity and release pro-inflammatory signals that reshape immune responses^125^. As the immune system regulates infection, tissue repair, inflammation, and cancer surveillance, its dysfunction contributes to organism-wide aging^126–128^. Conversely, tissue-level immune activity —including inflammation or loss of tolerance— may leave detectable traces in blood, offering a minimally invasive window into otherwise inaccessible compartments^129^. For instance, the female-specific expansion of three cytotoxic CD8+ TEM subpopulations (*GZMB+* TEM, NKT-like, and Temra), all expressing *GZMB* and exhibiting an NK-like innate cytotoxic feature^61^, suggests a systemic increase in cytotoxic potential **(Fig. 2D,E, Fig. S2D–F)**. Indeed, similar cytotoxic shifts have been reported in female endometrial CD8+ T cells after menopause due to hormonal withdrawal that lift suppression of perforin and granzyme expression^130–133^. Thus, cytotoxic signatures in immune circulating cells might echo changes in mucosal or barrier tissues. Beyond T cells, we find a female-specific expansion of CD14+ and CD16+ monocytes enriched for IFN-stimulated and inflammatory genes **(Fig. 2F)**, in line with previous findings linking these subsets to inflammaging^134,135^ and to impaired monocyte phagocytic function in older females^136^. Monocyte-driven inflammation has also been implicated in intestinal epithelial dysfunction in aging females^64^, reinforcing the idea that circulating immune changes may both signal and contribute to tissue-level immune decline. Together, these findings highlight that sex-specific immune trajectories in blood may serve as mechanistic indicators of systemic immune decline and, given the accessibility of blood sampling, also as practical biomarkers of aging and disease risk.

Our study has certain inherent limitations that should guide future research. First, expanding the cohort to include individuals from diverse ancestries and a wider age range, including pediatric and supercentenarian samples, would help capture the full spectrum and heterogeneity of immune aging. Second, although our analysis points towards non-linear immune aging trajectories, more robust statistical models and larger sample sizes are needed to fully capture these complex, sex-and cell type-specific patterns. Third, while we rely on a population cohort which excludes individuals with active infections, we lacked information on factors such as cytomegalovirus exposure, smoking, and comorbidities^137–140^, which may influence immune aging. Lastly, we focused on chronological aging, which may not fully reflect the biological age ^141–143^. Therefore, future studies incorporating additional layers, such as methylation profiles^144–146^ or telomere length^147^, could offer a more accurate proxy for biological age and provide deeper insight into immunosenescence. Despite these limitations, our work demonstrates the power of single-cell resolution to disentangle the complex interplay between aging, immunity, and sex. We show that stratifying analyses by sex uncovers key sex-specific features of immunosenescence, which, in some instances, have been misinterpreted as shared effects across sexes. This underscores the importance of treating sex as a source of biological variation —rather than a confounder— to avoid overgeneralization and ensure biologically accurate conclusions^30^. Ultimately, our findings lay the foundation for sex-tailored strategies to monitor immune aging and mitigate the burden of age-related immune dysfunction.

## Methods

### scRNA-seq dataset

#### Data overview and pre-processing

We analyzed scRNA-seq data over one million PBMCs (1,272,518 cells) from the OneK1K cohort^46^. This population cohort consists of 982 individuals (416 (42%) male and 566 (58%) female) of Northern European ancestry recruited in Hobart, Australia. The age of participants ranged from 19 to 97, with 73% of the participants being older than 60 years old. Participants with active immune diseases and non-European individuals were excluded from the study, following ancestry inference based on genotype data. The pre-processing of the scRNA-seq data (sample preparation and pooling, sequencing, quantification and alignment, demultiplexing, quality control and normalization) was conducted according to the original publication^46^.

#### Cell type annotation

We performed automated cell type classification using Azimuth to annotate the cells^51^. In detail, we conducted a supervised analysis guided by a reference dataset to enumerate cell types that would be challenging to define with an unsupervised approach. We mapped our scRNA-seq query dataset to a recently published CITE-seq scRNA-seq reference dataset of 162,000 PBMCs measured with 228 antibodies^51^. For this process, we used a set of specific functions from the R package Seurat (v4.0.0)^51,52^. First, we normalized the reference dataset using the Seurat::SCTransform function. Next, we found anchors between reference and query using a precomputed supervised PCA (spca) transformation with 50 dimensions using the Seurat::FindTransferAnchors function. We then transferred cell type labels from the reference to the query by projecting the query set on the UMAP embeddings of the reference using the Seurat::MapQuery function. To optimize computational resources, these steps were executed independently for each sequencing pool. Subsequently, the resulting cell type annotations were mapped back onto the complete dataset. We ended up with a total of 31 circulating immune cell subpopulations (https://github.com/powellgenomicslab/onek1k_scd). Finally, we filtered the dataset to retain only cell types with at least 5 cells in a minimum of 4 individuals, resulting in 24 distinct cell types **(Supplementary Table 1)**.

#### Sex-stratified cellular compositional analysis with age

To assess sex-specific differences in cell type proportions with age, we first calculated the relative proportions of each cell type within each sex. We then applied a centered log-ratio (CLR) transformation^148^ to account for the compositional nature of the scRNA-seq data. Specifically, for each donor, we divided the cell type proportions by their geometric mean, which converted the relative proportions into unconstrained values suitable for analysis with standard statistical methods.

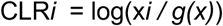

where *xi* is the cell type proportion being tested in donor *i*, and *g(x)* is the weighted geometric mean.

Lastly, for each cell type, we fitted a linear mixed model using the lmer function from the R package lmerTest (v3.1.3). We corrected for multiple testing using FDR, through the Benjamini–Hochberg method. We considered cell type proportions significantly changing with age at FDR < 0.05. The LMM used was:

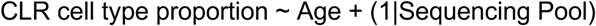

### Cell type abundance trajectory clustering analysis by sex

To identify aging-associated cellular dynamics, we grouped donors into 5-year age bins, stratified by sex. For each immune cell type and age bin, we computed the mean cell frequency and normalized these values using z-scoring across bins. Age bins with ≤5 donors (15-20 and 90-100), as well as cell types (CD4 CTL, NK_CD56bright, Plasmablast, NK Proliferating, dnT, pDC) with fewer than 5 donors in any bin, were excluded from analysis. We used these z-scored mean cell type frequencies to perform hierarchical clustering using the pheatmap function from the R package pheatmap (v1.0.12)^149^ with default parameters. Clusters of cell type-sex combinations were grouped based on similarity in their age-related abundance trajectories, and clusters were manually annotated into four trajectory categories: gradual decrease, inverted U-shape, gradual increase, and transition. Cell types were considered to exhibit sex-specific dynamics if male and female trajectories clustered into distinct groups.

### Cell type agnostic DA analysis with age

We used the R package miloR (v1.99.16, devel branch)^58^, a method designed to detect DA cell populations between conditions by modeling cell counts in neighborhoods of a KNN graph. Milo implements generalized linear mixed model (GLMM) for DA testing, which comes with computational and memory constraints. Due to these limitations, we could not run this analysis on the entire dataset and, instead, we randomly downsampled our data to 25% (316,939 cells out of a total of 1,272,489 cells). To ensure reproducibility, we repeated the downsampling three times and observed consistent results across replicates **(Fig. S5A,B, Supplementary Table 7)**. For the differential abundance analysis by sex, we split a downsampled dataset into female (369,888 of 739,776 total cells) and male (255,507 of 511,014 total cells) subsets.

#### Identification of neighborhoods of cells

To begin, we applied the batch-correction R package harmony (v1.0.3)^150^ to remove batch effects (i.e., sequencing pool) in our dataset. We used the function harmony::RunHarmony with default parameters and group.by.vars=”Sequencing Pool”. We then used the harmony-corrected PCA embedding to construct the KNN graph with the miloR::bulidGraph function, setting parameters k=100 and d=30. This function calculates the Euclidean distance between each cell and its KNN in the principal component space. We set a *k* parameter (*k*=100) that reduces the overall number of neighborhoods while keeping a relative small neighborhood size to achieve high resolution **(Fig. S5C,D)**. We also checked the count distributions for each neighborhood based on our variable of interest (age) using miloR::checkSeparation function with default parameters at several values of *k* **(Fig. S5E)**. This function determines whether neighborhoods exhibit perfect separation into all-zero versus non-zero counts for age, which can cause model failure in GLMM. The function outputs a logical classification (‘TRUE’ or ‘FALSE’) for each neighborhood, indicating the presence or absence of perfect separation, respectively. Identifying such neighborhoods is crucial for informing decisions on neighborhood subsetting or the recomputation of the nearest-neighbor graph to refine neighborhood definitions and ensure robust GLMM analysis. The distribution was similar among all *k* parameters. Next, we used the miloR::makeNhoods function to create neighborhoods of cells with parameters prop=0.05, k=100, and d=30, along with the graph-based refinement method (refinement_scheme=“graph”), which has been demonstrated to speed up this procedure^58^. Finally, we applied the miloR::countCells and miloR::calcNhoodExpression functions with default parameters to count the number of cells and their gene expression within each neighborhood. The same procedure was followed in the sex-stratified data (**Fig. S5F-H)**.

#### Testing for age-DA

We performed DA analysis with age for each sex using the miloR::testNhoods function with the following parameters: fdr.weighting=“graph-overlap”, glmm.solver=“HE-NNLS” and norm.method=“TMM”. The model was the following:

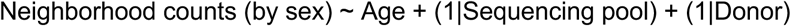

To control for multiple testing, we used an adapted version of a weighted FDR procedure^151^ for a KNN graph. In detail, this procedure accounts for the spatial overlap of neighborhoods, building upon an approach initially introduced in Cydar^152^. We employed an alternative weighting scheme to avoid distance calculations using the number of overlapping cells between each neighborhood^58^.

We then ran the function miloR::buildNhoodGraph with default parameters to create an abstracted graph of neighborhoods for visualization and miloR::annotateNhoods with coldata_col=“cell_type” to assign a cell type label to each neighborhood (here referred to as ‘Nhood cell type label’) based on the most frequent label of the neighborhood. For visualization purposes, changes in neighborhood abundance grouping by cell type, we omitted neighborhoods with a frequency of the major cell type label < 70%.

#### Identification of gene markers for age-DA enriched or depleted neighborhoods in males and females

To identify gene markers of neighborhoods significantly enriched or depleted with age in our sex-stratified DA within a specific cell type, we first extracted the expression data for all cells within each neighborhood of that cell type. In this case, we used the cell type annotation corresponding to the ‘Nhood cell type label’ obtained using the Milo framework as explained in a previous section. Then, we extracted the top 2,000 highly variable genes with Seurat::FindVariableFeatures function with default parameters. We used this set of genes to run Seurat::FindMarkers to identify significantly age-enriched or-depleted neighborhoods markers. Specifically, to find gene markers of age-DA enriched neighborhoods, we set *idents.1* to include all cells from neighborhoods that were significantly enriched with age (age-logFC > 0 & spatial-FDR < 0.05) and *idents.2* to include all cells from depleted neighborhoods with age, including both significant and non-significant ones (age-logFC < 0). Conversely, to identify gene markers of age-DA depleted neighborhoods, we set *idents.1* to include all cells from significantly depleted neighborhoods (age-logFC < 0 & spatial-FDR < 0.05), and *idents.2* to include all cells from enriched neighborhoods, including both significant and non-significant ones (age-logFC > 0). Any cells that were present in both *idents.1* and *idents.2* categories, due to the overlapping nature of the Milo graph, were excluded from the analysis. Genes were considered markers if they met the criteria of FDR < 0.05 and logFC > 1.

#### Clustering of age-enriched CD8 TEM neighborhoods

We performed unsupervised clustering of cells using a graph-based approach implemented in Seurat. Specifically, we constructed a K-nearest neighbor (KNN) graph based on Euclidean distances in PCA space and refined edge weights using Jaccard similarity (FindNeighbors function). Clustering was then performed via modularity optimization (e.g., Louvain algorithm) using the FindClusters function.

### Sex-stratified BCR repertoire with age in an independent longitudinal cohort^32^

We used previously processed BCR-seq data on PBMCs from a short-term longitudinal study^32^. In detail, donors were split into five groups according to their age, falling in 10-year intervals. Each age group followed over the course of 3 years with annual blood collection, which results in a total of 317 samples, with 2 technical replicates each one, from 166 healthy individuals aged 25-85 years (mean 49.38 ± 16.86) (https://www.synapse.org/Synapse:syn50268656). Briefly, the *Cell Ranger VDJ pipeline* (v6.1.2)^153^ was used to align BCR-seq reads to the GRCh38 genome and assemble BCR sequences, selecting those with one productive light and heavy chain pair for further analysis. Cell Ranger data were processed with the *Immcantation* framework (docker immcantation/suite:4.3.0) to re-assign V(D)J genes, define clonotypes using similarity thresholds based on sequence Hamming distances using DefineClones.py function from the Python package Change-O (v1.2.0) <u>(Gupta 2015)</u>, and calculate Gini coefficients using Gini function from the R package DescTools (v0.99.38) <u>(Signorell 2017)</u>. We then modeled clonal counts, separately for each sex, as a function of age, accounting for technical variables (number of cells (nCells) per donor, Donor_id, File_name, and Tube_id), using the following linear mixed model, using the lmer function from the R package lmerTest (v3.1.3):

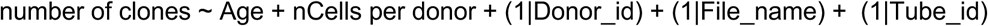

Due to a high sex imbalance in this dataset (130 males and 36 females), we considered downsampling male donors to match the number of female donors. However, we found that the observed age-associated signal was largely driven by a few male donors. As a result, downsampling would have masked meaningful biological variation, and we opted to retain the full dataset.

### Sex-stratified differential gene expression analysis with age

To identify DEGs with age within each of the 24 immune cell subpopulations, we performed sex-stratified differential expression analysis with age, conducting pseudobulk aggregation, pre-processing, and modeling separately within the males and females. First, we aggregated gene counts per donor to account for both zero-inflation and within-sample correlation, using the aggregateToPseudoBulk function from the R package dreamlet (v1.1.19)^154^ with default parameters. Second, we applied sample-and gene-level filters on the pseudobulk counts to ensure data quality and statistical robustness, using the processAssays function from the R package dreamlet (v1.1.19)^154^ with default parameters. Specifically, within each cell cluster, we retained samples with at least 5 cells (min.cells = 5) and kept only cell clusters with a minimum of 4 such samples (min.samples = 4). Genes were retained if they have ≥ 5 reads (min.count = 5) in at least 40% of the samples (min.prop = 0.4). Genes falling these criteria were excluded due to insufficient power for differential expression analysis and to reduce the multiple testing burden. Third, we applied an inverse-normal rank-transformation to the log2CPM expression values of each gene —referred to here as ‘pseudobulked normalized expression’— following the approach of ^155^. Briefly, for each gene within each immune cell subpopulation, we ranked the pseudobulk expression values of all individuals, assigned random ranks for ties, and replaced each observation with the corresponding quantile from a normal distribution with the same mean and standard deviation as the original expression data. This transformation reduces the impact of zero-inflation and outliers while preserving the scale of the original expression values. Fourth, we fit a linear mixed model for each gene to assess associations between gene expression and age (modeled as a continuous variable), while controlling for other sources of technical variation (sequencing pool and nCells per donor), using the lmer function from the R package lmerTest (v3.1.3). We corrected for multiple testing using FDR, through the Benjamini–Hochberg method. The model used was:

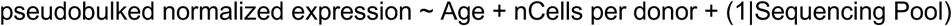

Genes were considered differentially expressed when the age coefficient was significant at an FDR < 0.05.

To identify sex-specific age gene expression effects, we additionally fitted an alternative model including an age-by-sex interaction term, using the same pre-processing pipeline (steps 1–3). We corrected for multiple testing using FDR, through the Benjamini–Hochberg method. The model used was:

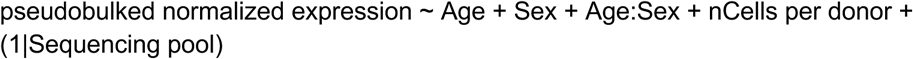

Genes were considered differentially expressed when the interaction term was significant at a nominal p-value < 0.01.

### Downsampling analysis for the sex-stratified differential gene expression analysis with age

#### Downsampling to the same number of donors per cell type between sexes

The number of male and female donors differed across the 24 immune cell subpopulations, with male donors consistently underrepresented **(Fig. S1C)**. To assess whether differences in sample size between sexes influenced the detection of age-associated DEGs, we performed a downsampling analysis. For each cell type, we randomly subsampled female donors to match the number of male donors, repeating this process four times to account for sampling variability. Each replicate was processed using the same pseudobulk aggregation, preprocessing, and modeling pipeline as described in the main analysis.

#### Downsampling to the cell type with minimum number of donors within sexes

The 24 immune cell subpopulations were detected in different numbers of donors **(Fig. S1C)**. To assess whether differences in sample size between cell types affected the detection of age-associated DEGs within each sex, we performed a downsampling analysis. For each cell type and sex, we randomly subsampled donors to match the number of donors in the cell type with the lowest sample size that still yielded at least one age-associated DEG (i.e., 468 donors in female gdT cells and 347 donors in male CD4+ CTL cells) **(Fig. 3A)**. This downsampling was repeated ten times per cell type to account for sampling variability. Each replicate was processed using the same pseudobulk aggregation, preprocessing, and modeling pipeline as described in the main analysis.

### Sex-stratified differential gene expression analysis with age across three age groups (early, mid, and late)

To investigate whether gene expression changes with age follow different patterns across the adult lifespan, we stratified individuals into three overlapping age groups: early (<50 years), mid (40–60 years), and late (≥50 years). Within each age group and sex, we performed differential gene expression analysis with age using the same pseudobulk aggregation, preprocessing, and modeling pipeline as described in the main analysis.

To assess the potential impact of sample size differences across age groups **(Fig. S4A)**, we performed a downsampling analysis. For each cell type and sex, we randomly subsampled donors to match the number of donors in the age group with the smallest sample size (i.e., 111 donors in the early group (<50 years old) for females and 80 donors in the mid group (40–60 years old) for males). This downsampling was repeated 4 times per cell type to account for sampling variability. Each replicate was processed using the same pseudobulk aggregation, preprocessing, and modeling pipeline described above.

### Sex-stratified breakpoint differential gene expression analysis with age

To identify temporally localized shifts in immune gene expression —referred to as “breakpoints” or “waves”— we tested, for each immune cell subpopulation and sex, whether short age intervals were characterized by significant changes in gene expression compared to adjacent time periods. These breakpoints reflect systemic chronological signatures that may indicate periods of rapid molecular change. To this end, we reimplemented the algorithm described in^71^ to uncover waves of gene expression changes across the lifespan in a sex-stratified manner. For each cell type and sex, we subsampled the data by iteratively applying a 30-year sliding window centered on a series of unique ages *l*_k_ spanning the sampled age range (*k*_min_ to *k*_max_). Within each window, we performed differential gene expression analysis by comparing donors in the preceding 15-year age interval to those in the following 15-year interval (e.g., 35–50 vs. 50–65 at a window centered on age 50). The window center was shifted one year at a time, generating a continuous scan across the adult lifespan. To mitigate the impact of outliers, variations in sample density across ages and temporal fluctuations, we restricted our analysis to window centers between 34 and 82 years old. This range was chosen by extending the minimum and maximum ages by ±15 years to ensure adequate representation across age groups. We further required each window center to include at least 130 age-balanced donors, with a minimum of 20% representation from each age group to preserve age balance.

Each subsampled dataset was processed using the same pseudobulk aggregation, preprocessing and modeling pipeline as described in the main analysis. In this case, age was modeled as a binary categorical variable (young [*l*_k_ - 15yo; *l*_k_] vs. old [*l*_k_; *l*_k_ + 15yo]), using the following model:

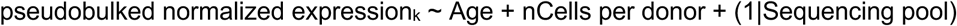

We corrected for multiple testing using FDR, through the Benjamini–Hochberg method. Genes were considered differentially expressed at a given window center when the age coefficient was significant at an FDR < 0.1. To assess the robustness of the breakpoint detection, we repeated the analysis using alternative sliding window widths of 10, 20, and 40 years.

### Inferring gene expression chronological aging trends by sex

To estimate gene expression trajectories across the lifespan, we first z-scored gene expression counts for each cell type and sex. A linear mixed model was then fitted for each gene using the lmer function from the R package lmerTest (v3.1.3) to regress out batch effects associated with sequencing pool and nCells per donor, consistent with the differential gene expression analysis. Residuals from these models were used to fit locally estimated scatterplot smoothing (LOESS) regression using the loess function from the R package stats (v4.3.0). To group genes with similar aging trajectories, we computed pairwise Euclidean distances between LOESS estimates using the dist function (method = “euclidean”), followed by hierarchical clustering using the hclust function (method = “complete”) from the same package. Finally, to interpret the biological relevance of each gene cluster, we performed functional enrichment analysis querying the Gene Ontology (GO) database^156,157^ as described below. This analysis was constrained to genes that were significantly age-associated in the “breakpoint” analysis.

### Functional enrichment analysis

#### Over-representation analysis (ORA)

We performed ORA using the R package clusterProfiler (v4.12.0)^158^ throughout the manuscript. We ran the analysis against the GO database^156,157^ using clusterProfiler::enrichGO function, with default parameters except foront=“GO”, which was set to “ALL” only when analyzing cell type-specific DEGs. For cell type-specific ORA, the background set comprised all genes expressed in the corresponding cell type. When analyzing age-DEGs shared across multiple cell types, the background was defined as the intersection of expressed genes across those cell types. Multiple testing correction was applied using the Benjamini–Hochberg method to control the FDR, and terms with FDR < 0.05 were considered significant. In some cases **(Fig. 2C**, **Fig. 3B, Fig. S3F, Fig. S4G)**, for visualization purposes, we generated non-redundant sets of GO terms. To do so, we used the R package rrvgo (v1.16.0)^159^, which simplify the redundancy of GO sets by grouping similar terms based on their semantic similarity. We first applied the rrvgo::calculateSimMatrix function to get the similarity matrix between terms, and then, the rrvgo::reduceSimMatrix function to group terms, setting a similarity threshold to collapse redundant terms. Within each group, the term with the highest uniqueness score was selected as the representative. For comparisons spanning multiple cell types, enrichment results were merged across cell types before applying term reduction.

In addition, we conducted ORA using the DisGeNET database^74,75^, using the disease_enrichment function from the R package disgenet2R (v1.2.2)^74,75^, with default parameters. As explained above, FDR was controlled using the Benjamini–Hochberg method, and terms with FDR < 0.05 were considered significant.

#### AUCell Gene Set Enrichment Analysis (GSEA) using autoimmune, interferon, and inflammatory gene sets

We performed single-cell GSEA using the R package escape (v1.99.1)^160^, which integrates multiple GSEA approaches optimized for scRNA-seq data. We used the escape::runEscape function, with default parameters, except for min.size=0, groups=5000, method=“AUCell”. AUCell^161^ calculates the area under the curve (AUC) to determine whether specific gene sets are enriched among the top 10% expressed genes in each cell, yielding a per-cell enrichment score. To account for sparsity and dropout in single-cell data, we normalized the AUCell scores using the escape::performNormalization function, which ensures all enrichment values are positive and suitable for downstream modeling^160^.

Then, within each sex, we fitted a GLMM with a Gaussian distribution for each gene set, using the glmmTMB function from the R package glmmTMB (v1.1.9), to assess associations between enrichment scores and age (modeled as a continuous variable) across different cell types. We included random intercepts for sequencing pool and donor to account for technical and inter-individual variability, respectively. We corrected for FDR through the Benjamini–Hochberg method. The model used was:

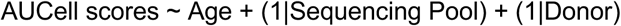

Associations between gene set enrichment scores and age were considered significant at FDR < 0.05.

### Interferon and inflammatory enrichments of age-enriched CD14+ Monocytes in females

To evaluate interferon-stimulated and inflammatory gene set enrichment in CD14+ Monocyte neighborhoods, we followed the same approach as in^155^. Briefly, we defined our interferon-stimulated gene set (n=182) by selecting the genes included in the union of GSEA hallmark database (https://www.gsea-msigdb.org/gsea/msigdb/genesets.jsp?collection=H) IFN-α response (‘HALLMARK_INTERFERON_ALPHA_RESPONSE’, n=97) and IFN-γ response (‘HALLMARK_INTERFERON_GAMMA_RESPONSE’, n=200) gene sets, but excluded those from the inflammatory response set (‘HALLMARK_INFLAMMATORY_RESPONSE’, n=200). Similarly, we defined our inflammatory gene set (n=158) by selecting the genes included in the inflammatory response set, but excluded those from the IFN-α and IFN-γ response gene sets **(Supplementary Table S3).** For visualization purposes **(Fig. 2F),** we computed the mean AUCell score among neighborhoods with at least 15 cells. We then compared female age-enriched and male age-depleted DA CD14⁺ monocyte neighborhoods using a Wilcoxon rank-sum test, implemented via the wilcox.test function from the R package stats (v4.3.0).

### Autoimmunity score enrichments in CD4+ TCM

To assess enrichment for autoimmunity in CD4+ TCM we used a curated autoimmune gene set from previous studies^79,80^. For visualization purposes **(Fig. 4B),** we computed the mean AUCell score per donor. We then compared scores between sexes using a Wilcoxon rank-sum test implemented via the wilcox.test function from the R package stats (v4.3.0), stratifying donors into two age groups: younger than 50 years and older than 50 years.

#### GSEA for immune subpopulation signature marker genes

We used the fgsea R package^162^ to evaluate whether neighborhood marker genes were enriched in previously defined immune subpopulation gene signature from^32^. Genes were ranked by their average logFC, and enrichment was evaluated for the following Milo-identified neighborhood groups: *i)*CD8+ TEM neighborhoods in aging females: Marker genes from age-enriched CD8+ TEM neighborhoods were tested against signature genes representing CD8+ TEM subpopulations, including NKT-like, Temra, *GZMB+* CD8+ TEM, and *GZMK+* CD8+ TEM. *ii)* B naive and intermediate cell neighborhoods in aging males: Marker genes from age-enriched B naive and intermediate cell neighborhoods were tested against signature gene for B cell subpopulations, specifically CD5+ atypical memory B cells and activated B cells. *iii)* CD4+ TCM neighborhoods in aging females: Marker genes from both age-enriched and age-depleted CD4+ TCM neighborhoods were tested against signature genes of CD4+ T helper subpopulations, including Th22, Th17, Th1/Th17, and Th1 cells.

### Overlap of sex-stratified age-DEGs with GWAS hits

The GWAS data was obtained from the GWAS Catalog (https://www.ebi.ac.uk/gwas/). We identified genes associated with the strongest SNPs using the “MAPPED GENE(S)” column, including those located within or near these SNPs. Then, per each cell type, we assessed whether sex-stratified age-DEGs were enriched for these disease-associated genes using the fisher.test function from the R package stats (v4.3.0) with default parameters. We applied the Benjamini–Hochberg method to adjust for FDR, considering a cell type to be significantly enriched with age in a particular gene set at FDR < 0.05.

### Sex-stratified cellular compositional and differential gene expression analyses with age on manually curated subsets of CD8+ TEM, CD14+ Monocytes and B naive cells

To further characterize the sex-specific CD8+ TEM, CD14+ Monocyte, and B naive subpopulations that change with age in our cell type–agnostic age-DA analysis, we classified these cell types using top marker genes identified in the previous analysis (**Supplementary Table 3**) and supported by the literature^32,36,56,163,164^. Specifically, we used *GZMB* for CD8+ TEM, *ISG15* for CD14+ Monocytes, and *ABCA6* for B cells. Thus, within each cell type, cells were then divided into two subpopulations based on marker gene expression: marker-positive (≥1 count) or marker-negative (<1 count) (*GZMB+* CD8+TEM and *GZMB*-CD8+TEM; *ISG15*+ CD14+ Mono and *ISG15*-CD14+ Mono; *ABCA6*+ *B cells* and *ABCA6*-*B cells*). We then performed sex-stratified differential gene expression analysis with age within each subpopulation, following our previously described approach. Additionally, for CD8+ TEM, we conducted a sex-stratified composition analysis based on *GZMB*-defined subtypes to assess age-related shifts.

### Replication of sex-stratified cell type abundance changes with age with external datasets

To validate our main findings from the OneK1K cohort^46^ (discovery dataset), we relied on two external datasets^62,32^ (replication datasets) of healthy individuals spanning a wide age range. While these two datasets provided valuable replication potential, they differed in sample size and sex distribution compared to the discovery cohort. To accommodate these differences, we adjusted statistical significance thresholds to ensure robust inference where necessary.

#### Oelen et al 2022^62^

We used a subset of previously processed scRNA-seq data^62^ on unstimulated PBMCs from 72 healthy LifeLines Deep donors aged 27-77 years (mean 56.83 ± 12.7) (https://www.eqtlgen.org/sc/datasets/1m-scbloodnl-dataset.html). In short, scRNA-seq data was generated using the 10X Genomics Chromium Single Cell 3’ v2 chemistry and libraries were 150 bp paired-end sequenced on Illumina’s NovaSeq6000. The Cellranger v3.0.2 pipeline was used with default parameters to demultiplex, generate FASTQ files, map reads to the GRCh37 reference genome and generate a unique molecular identifier (UMI)-corrected count matrix per cell. To assign cells to one of the eight individuals in a lane, Demuxlet was used. After quality control, 54,373 cells remained. To enable a direct comparison between the two datasets, we employed the same cell annotation strategy used for the OneK1K cohort. In detail, we performed automated cell type classification using Azimuth to annotate the cells^51^. We mapped our scRNA-seq query dataset to a recently published CITE-seq scRNA-seq reference dataset of 162,000 PBMCs measured with 228 antibodies^51^. For this process, we used a set of specific functions from the R package Seurat (v4.0.0)^51,52^. First, we normalized the reference dataset using the Seurat::SCTransform function. Next, we found anchors between reference and query using a precomputed supervised PCA (spca) transformation with 50 dimensions using the Seurat::FindTransferAnchors function. We then transferred cell type labels from the reference to the query by projecting the query set on the UMAP embeddings of the reference using the Seurat::MapQuery function. We ended up with a total of 31 circulating immune cell subpopulations. Finally, we filtered the dataset to retain only the 24 cell types present in the OneK1K data after applying specific filters (i.e., at least 5 cells in a minimum of 4 individuals, **Supplementary Table 1**). We used this dataset to replicate the sex-specific abundance changes with age **(Supplementary Table 3)**. To do so, we applied the same pipelines as in the OneK1K data.

#### Terekhova et al 2023^32^

We used previously processed scRNA-seq data^32^ on PBMCs from a short-term longitudinal study. In detail, donors were split into five groups according to their age, falling in 10-year intervals. Each age group followed over the course of 3 years with annual blood collection, which results in a total of 317 samples, with 2 technical replicates each one, from 166 healthy individuals aged 25-85 years (mean 49.38 ± 16.86) (https://www.synapse.org/Synapse:syn50542388). Briefly, scRNA-seq data was obtained using the 10x Genomics Chromium Single Cell 5’ v2 chemistry and libraries were sequenced on a NovaSeq S4 flow cell (300 cycles), targeting 25,000 read pairs per cell. In total, 107 sequencing outputs were generated from the 317 samples. Each output was then independently demultiplexed using a combined hashtag-based and genotype-based demultiplexing strategy, including Seurat v4.0.5 *HTODemux* and *minimap2* v2.17-r941, *freebayes* v1.3.2, and *souporcell* v2.5. This strategy improved demultiplexing efficiency and accuracy compared to using a single method. After quality control, 1,916,367 cells remained. We used this dataset to replicate the sex-specific abundance changes with age **(Supplementary Table 3)**. To do so, we applied the same pipelines as in the OneK1K data.

## Data and code availability

The original public-access data from^46^, including OneK1K single-cell gene expression and genotype data are available via Gene Expression Omnibus (GSE196830). The cell by gene data are available at Human Cell Atlas (HCA) (https://cellxgene.cziscience.com/collections/dde06e0f-ab3b-46be-96a2-a8082383c4a1). All the custom scripts developed for this study have been made openly accessible and can be found on the GitHub repository at https://github.com/mariasr3/2024_SingleCell_ImmuneAging. Processed data files derived from the analysis conducted in this paper are deposited at Zenodo: https://doi.org/10.5281/zenodo.14035325. By visiting this repository, researchers and interested individuals can access and use the custom scripts for their own analysis or to replicate the study’s findings. Supplementary tables can be found: https://drive.google.com/drive/folders/1owmMW9wjL3gemao0-jEYb_dpPrhRl7Dz?usp=drive_link

## Acknowledgements

M.S.R was supported by a predoctoral “AGAUR-FI Joan Oró” fellowship from “Secretaria d’Universitats i Recerca del Departament de Recerca i Universitats de la Generaliat de Catalunya i del Fons Europeu Social Plus” (2024 FI-3 0065). A.R.C was supported by a predoctoral fellowship “Formación Personal Investigador (FPI)” from “el Ministerio de Ciencia, Innovación y Universidades (MCIN)” and “la Agencia Estatal de Investigación (AEI)” (MCIN/AEI FPI with code PRE2019-090193). M.M. was supported by a grant PID2019-107937GA-I00 funded by MCIN/AEI/10.13039/501100011033 and a grant RYC-2017-22249 funded by MCIN/AEI/10.13039/501100011033 and by “ESF Investing in your future”. J.E.P is supported by the National Health and Medical Research Council Fellowship (APP1107599) and with the support of the Fok Family in memory of Dr. and Mrs. Wing Kan Fok. L.F. is supported by a grant from the Dutch Research Council (ZonMW-VICI 09150182010019), and a sponsored research collaboration with Biogen, Roche and Takeda. This work has been co-financed with the Catalan government grant 2021 SGR01577. This work is co-financed by Oncode Institute, which is partly funded by the Dutch Cancer Society. This project has received funding from the European Union’s Horizon Europe research and innovation programme under grant agreement No 101057553. M.G.P.W. is supported by a NWO Vidi grant of the Dutch Research Council (VI.Vidi.223.041). We thank Winona Oliveros, José Miguel Ramirez, and Fairlie Reese for their valuable feedback during the final stages of this work, and Victor Jiménez for his insights on B cell leukemias.

## Declaration of generative AI and AI-assisted technologies in the writing process

During the preparation of this work, the authors used ChatGPT to enhance the clarity and language of the text and Perplexity to locate relevant references for the manuscript. After using these tools, the authors reviewed and edited the content as needed and take full responsibility for the content of the publication.

## Author contribution

M.S.R. designed and performed all analysis, A.R.C designed, performed and supervised all analysis.

F.O. performed specific analysis (i.e., overlap of sex-stratified age-DEGs with GWAS hits).

S.B. and J.A.H. performed specific analysis (i.e., OneK1K cell type annotation).

R.O. performed specific analysis (i.e., cell type annotation of the subset of data from^62^ used for several replication analyses).

M.G.P.W, L.F, and J.P. provided overall feedback.

M.M. conceived the study and supervised all analysis. M.S.R., A.R.C, and M.M, wrote the manuscript.

All authors read and agreed on the contents of this manuscript.

## Supplementary Figures

**Figure S1.**
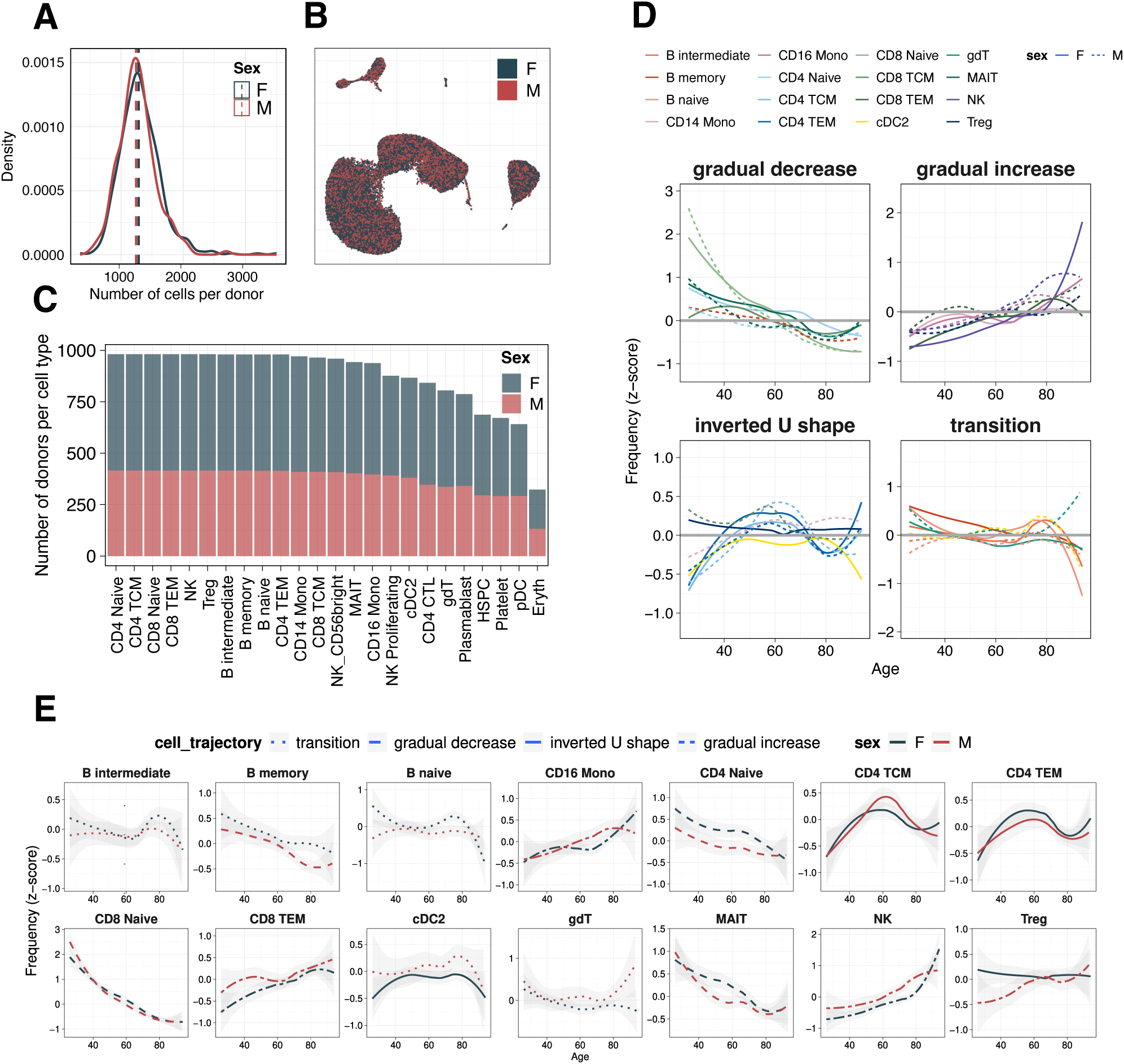
Data overview and immune cell trajectories across the lifespan. **A.** Distribution of the number of cells per donor, shown separately for males (red) and females (blue). **B.** UMAP plot of all immune cells, colored by sex. For visualization purposes we downsampled to 500,000 cells (representing ∼50% of the total cells) **C.** Number of donors per cell type and sex. **D.** Immune cell trajectory groups over the lifespan. The y-axis represents normalized cell frequency (z-score), and the x-axis represents age. Color indicates cell type and line type indicates sex. **E.** Immune cell trajectory per cell type over the lifespan. The y-axis represents normalized cell frequency (z-score), and the x-axis represents age. Color indicates sex and line type indicates trajectory group.

**Figure S2.**
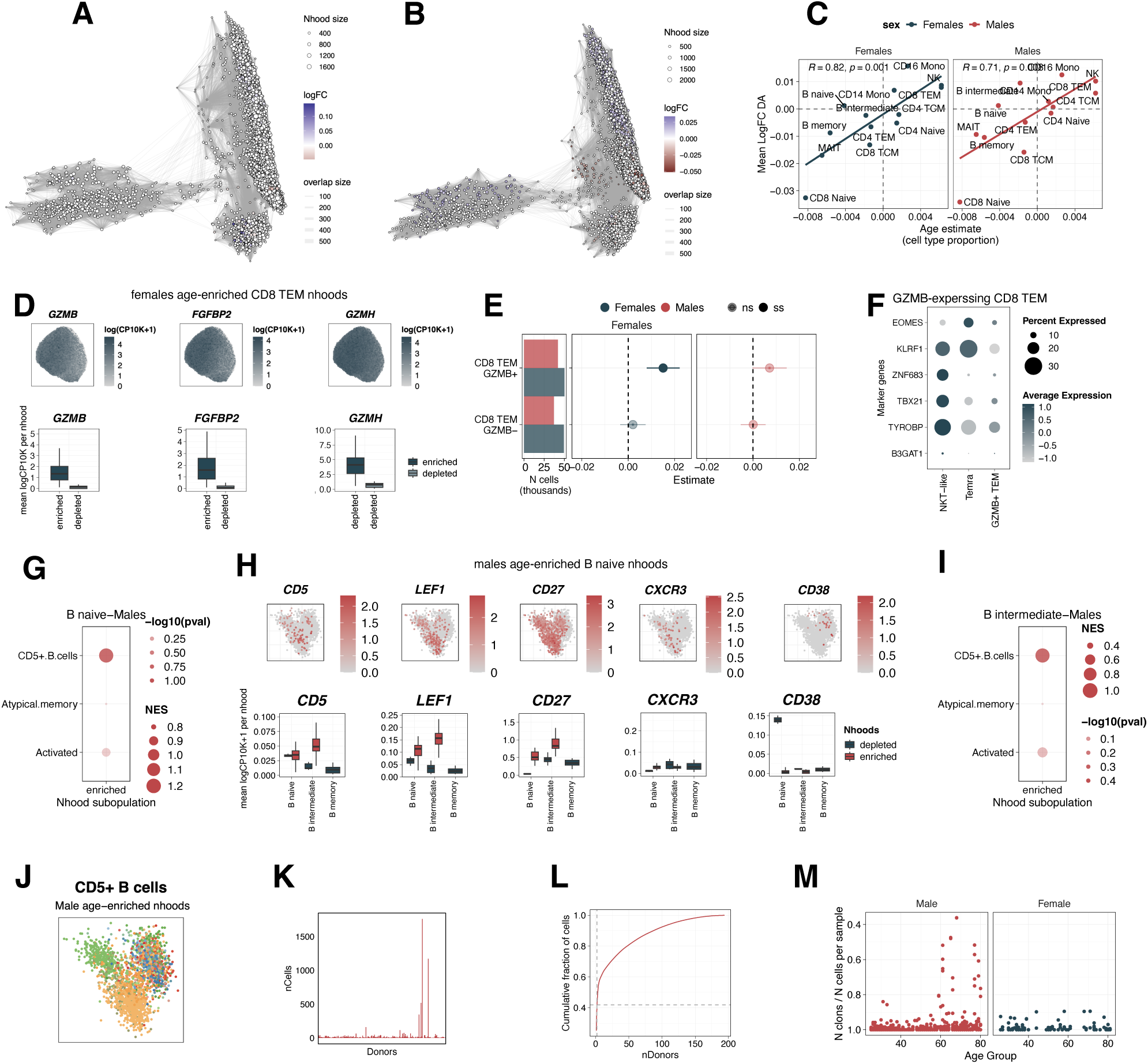
Sex-stratified differential abundance analysis. Nhood (neighborhood) graph of the results from differential abundance testing with age, separately by males **(A)** and females **(B).** Nodes represent Nhoods, colored when they are significant (Spatial-FDR < 0.05), and edges correspond to the number of cells shared between Nhoods. **C.** Correlation of the mean logFC per cell type across Nhoods (y-axis) versus the age-estimate of the cell type compositional analysis (x-axis). **D.** Top. UMAP plots of female-age-enriched CD8+ T effector memory (TEM) cells colored by the expression of the three top marker genes (*GZMB, FGFBP2*, and *GZMH*). Bottom. Average log-normalized, scaled expression per Nhood of the same genes. Only Nhoods that are significantly enriched or depleted with age are shown. **E.** Left. Total number of cells analyzed per CD8+ TEM subpopulation and sex. Center. Cell type composition analysis estimates with age from *GZMB*+/-CD8+ TEM subpopulations in females. Right. Cell type composition analysis estimates with age from CD8+ TEM subpopulations. **F.** Average expression of marker genes for *GZMB-*expressing cytotoxic subpopulations of CD8+ TEM. **G.** Gene set enrichment analysis (GSEA) of marker genes of age-enriched B naive Nhoods in males compared to transcriptional signatures of B naive subpopulations from <u>Terekhova et al., 2023</u>. Significant enrichments (FDR < 0.05) are highlighted. **H.** Top. UMAP plots of male-age-enriched B naive and intermediate cells, colored by the expression of the top marker genes. Bottom. Average log-normalized, scaled expression per Nhood in males of the same marker genes.Only Nhoods that are significantly enriched or depleted with age are shown. **I.** Same as panel **G** but for B intermediate cells. **J.** UMAP plot of male *CD5+* B subpopulation colored by donor. **K.** Number of cells per male donor belonging to *CD5+* subpopulation. Donors are sorted by increasing age. **L.** Cumulative fraction of cells from *CD5+* B subpopulation belonging to a particular donor. **M.** Ratio of number of clones by number of cells per sample in *CD5+* B cells through the lifespan in males (left) and females (right)

**Figure S3.**
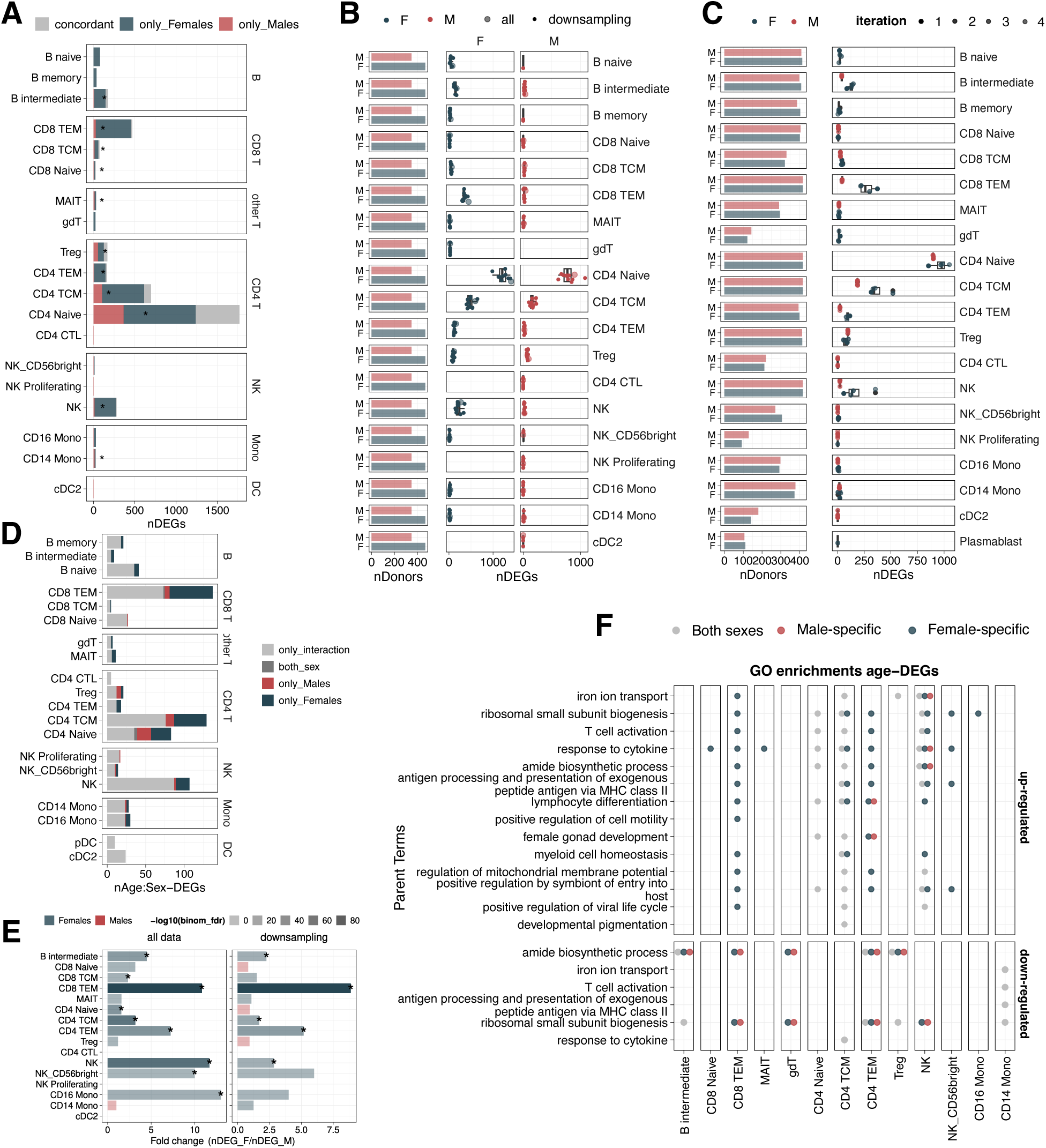
Sex-stratified linear age-differential expression analysis. **A.** Overlap of age-differentially expressed genes (age-DEGs) in each sex-stratified analysis. Asterisks indicate significant overlaps (FDR< 0.05). **B.** Left. Total number of donors analyzed per cell type. Right. Number of age-DEGs detected in males and females after downsampling donors to match the number of donors in the cell type with the lowest sample size that still yielded at least one age-associated DEG (i.e., 468 donors in female gdT cells and 347 donors in male CD4+ CTL cells). Light dots represent age-DEG counts from ten independent downsampling iterations, and the dark one correspond to the real number of age-DEGs. **C.** Same as panel **B** but downsampling to the same number of donors between sexes for each cell type. **D.** Number of DEGs (nominal p-value < 0.01) from a model including the Age:Sex interaction term, colored to indicate whether each gene was also detected in the sex-stratified age-DEA. **E.** Ratio of the number of age-DEGs in females versus males, colored by the sex with the higher number of age-DEGs. Asterisks indicate statistically significant sex bias (binomial test, FDR < 0.05). Intensity of the color corresponds to –log₁₀(FDR) value of the binomial test. Results are shown for the full dataset (left) and after downsampling to equal numbers of donors between sexes (right). **D.** Non-redundant Gene Ontology (GO) terms from enrichment analysis of upregulated (top) and downregulated (bottom) age-DEGs. Each dot indicates whether the corresponding GO term is enriched in sex-shared (grey), male-specific (red), or female-specific (blue) age-DEGs.

**Figure S4.**
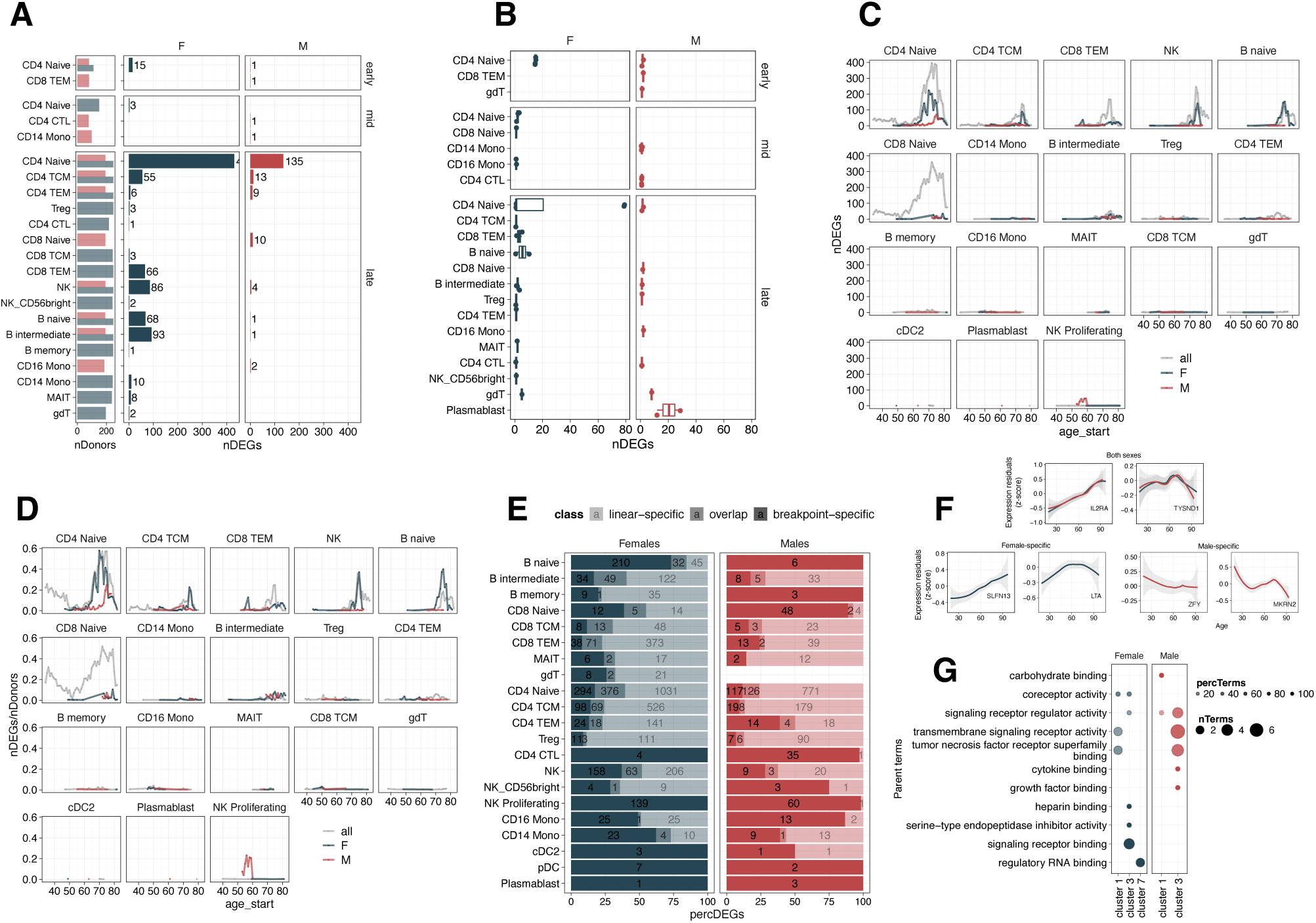
Sex-stratified non-linear age-differential expression analysis. **A.** Left. Number of donors per cell type. Right. Number of age-differentially expressed genes (age-DEGs) per cell type across age bins: early (<50 years old), mid (40–60 years old), and late (≥50 years old), separately in males and females. **B.** Number of age-DEGs detected in males and females after downsampling to match the number of donors in the smallest age group (early (<50 years old) in females —111 donors—, and mid (40–60 years old) in males —80 donors—). Each dot represents the result from one of the four independent downsampling iterations. **C.** Number of age-DEGs identified through breakpoint analysis in each cell type across the lifespan in females (blue), males (red) and the combined dataset (grey). **D.** Same as panel **C** but the y-axis shows the number of age-DEGs normalized by the number of donors in each comparison. **E.** Percentage of age-DEGs across cell types in each sex, identified trough: breakpoint analysis (breakpoint-specific), linear models (linear-specific) or both (overlap). **F.** Representative examples of gene expression trajectories illustrating linear (left) and non-linear (right) transcriptional changes in immune aging. Top: age-DEGs in both sexes. Bottom: Female-specific age-DEGs (left) and male-specific age-DEGs (right). **G.** Non-redundant Gene Ontology (GO)

**Figure S5.**
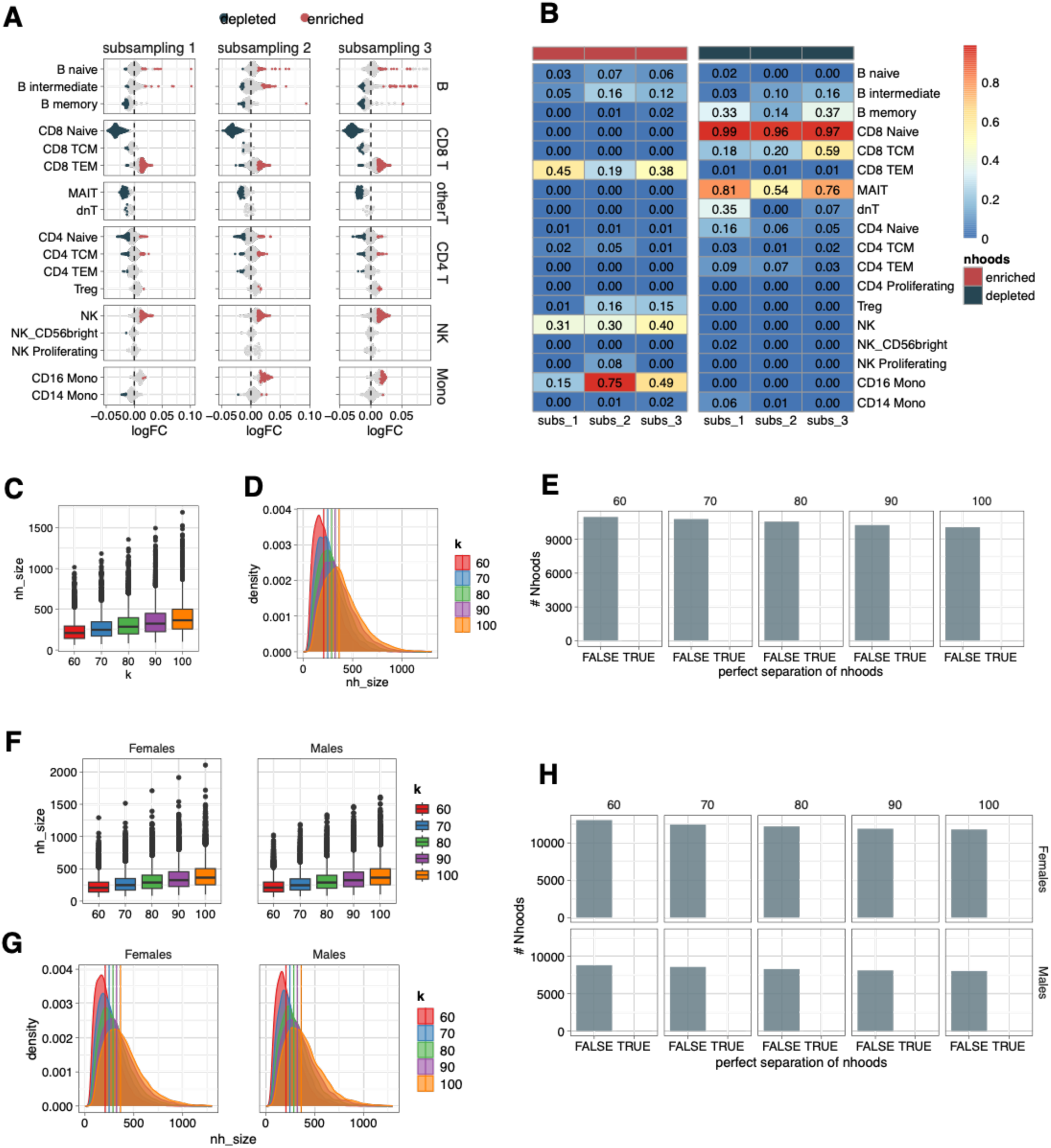
Differential abundance analysis: replication using data subsamplings and fine-tuning the *k* parameter for the KNN graph. **a.** Beeswarm plot of the distribution of neighborhood (Nhood) logFC with age for each data subsampling (25% of the total cells; 316,939 cells out of a total of 1,272,489 cells). Each dot corresponds to a Nhood of cells assigned to a dominant cell type (>80% of cells within the neighborhood (see **Methods**)). Age-DA Nhoods (Spatial-FDR < 0.05) are colored. **b.** Proportion of age-DA enriched (left) and depleted (right) Nhoods in each data subsampling. **C.** and **D.** To identify the proper value of nearest neighbors to consider for the graph building (*k*), we set different *k* values (60, 70, 80, 90 and, 100), and compared the distribution of the number of Nhoods for each Nhood size **(C)** and the distribution of the Nhood size across different values of *k* **(D)**. Vertical lines in **C** correspond to the mean Nhood size. **E.** Number of Nhoods that have perfect data separation based on age categories (TRUE) versus the ones that do not (FALSE) at different values of *k*. Age categories have been defined as: young (<20 years), middle-aged (21–40 years), and old (>41 years) (see **Methods**)).

